# Convergent genome- and gene-level constraints shape repeated environmental adaptation in grasses

**DOI:** 10.64898/2026.06.01.729361

**Authors:** Sheng-Kai Hsu, Aimee J Schulz, Charles O Hale, Germano Costa-Neto, Zachary R Miller, Michelle C Stitzer, Travis Wrightsman, Jingjing Zhai, Wei-Yun Lai, Henry D Dawson, Qifan Wang, Taylor M AuBuchon-Elder, Lara J Brindisi, Elad Oren, Yun Luo, Beatrice A Konadu, Jonathan O Ojeda-Rivera, Matt Pennell, Elizabeth A Kellogg, M Cinta Romay, Edward S Buckler

## Abstract

Grasses (Poaceae) dominate terrestrial ecosystems and sustain global food security, yet the genomic principles enabling their repeated adaptation to extreme environments remain unresolved. Combining dense phylogenomic sampling, global environmental data, and genomic large language models (gLLMs), we characterize the mutational targets underlying environmental shift across 687 genomes from 561 species spanning 17 climate zones. We identify 17-30 phylogenetically independent transitions into extreme temperature, water, and soil environments, accompanied by convergent shifts in genome-scale molecular properties, including the Nitrogen-to-Carbon balance and the biosynthetic cost of the proteome. Our gLLMs-informed phylogenetic mixed modeling framework identifies 330 genes that repeatedly associated with distinct axes of environmental variation, highlighting the importance of protein modification and localization within extracellular and organellar compartments. Overlaying independent convergent selection tests identifies 27 high-confidence candidates for further characterization. Together, our results show that grass adaptation is canalized by layered constraints at genome-wide and gene-specific scales, producing predictable evolutionary trajectories.

## Introduction

Originating approximately 100 million years ago in tropical Gondwana, Poaceae (grasses) has since diverged, dispersed, and converged across the globe through tectonic movements, climatic shifts, and biotic dispersal, becoming one of the largest and most ecologically successful plant families^1,2^. Today, Poaceae encompasses a total of 791 genera and ∼11,800 species, including the cereals that underpin global food security (e.g., maize, wheat, rice, sorghum, sugarcane)^3^. Grass species are found on roughly 40% of Earth’s land surface^4^, occupying diverse ecological niches ranging from the arid tropical savannas to the freezing arctic tundra^4–7^.

This ecological dominance is underpinned by several key trait innovations that evolved in concert with the family’s diversification. These include the repeated evolution of the annual life history strategy, which allows rapid reproduction in seasonal environments^8,9^; the C4 photosynthetic pathway, which evolved independently >20 times in grasses to boost water- and nitrogen-use efficiency in hot, low-CO2 and open habitats^10–12^; and photoperiod insensitivity, enabling expansion into high latitudes with short growing seasons^13^. Deciphering the genetic basis of these adaptations is thus crucial not only for explaining the evolutionary rise of this dominant plant family but also for engineering climate-resilient crops in an era of accelerating climate change.

The diverse environmental niches occupied by grasses and the major phenotypic innovations in the clade may involve convergent changes at multiple biological scales of the genome. At a broad scale, adaptation to contrasting environments is hypothesized to involve genome-wide molecular and physiochemical modifications. Across angiosperms, environmental selection has been proposed to constrain genome size, favoring smaller genomes in high-productivity, nutrient-limited, or metabolically demanding environments where smaller cells are required for faster cell division and higher photosynthetic rates^14–19^. Environmental pressures have also been hypothesized to shape GC content, potentially increasing DNA stability in extreme environments^18,20^, and to influence codon usage bias to optimize translational efficiency under stresses^21–23^. Simultaneously, selection can target specific, conserved stress-response pathways and genes. Comparative studies among dozens of grass species have successfully revealed key adaptive genes associated with specific abiotic factors. For instance, the C4 photosynthetic pathway has been well-characterized as a mechanism for enhancing water-use efficiency under drought and heat stress^11,24^. Similarly, the *VRN* (vernalization) regulatory network is critical for overwintering competence^25–28^, while the *CBF*/*DREB* pathway serves as a master regulator of tolerance to cold, freezing, drought and osmotic stresses^29–31^. Late embryogenesis abundant (LEA) proteins, particularly dehydrins, act as downstream effectors of cold and drought stress by stabilizing cellular structures during dehydration and freezing^32–34^, and their accumulation contributes to freezing tolerance in several grass species^33,35–37^.

Together, these observations suggest that environmental adaptation may involve coordinated variation across genome-wide features, sequence-level modifications, and pathway-specific responses. Although evidence for these mechanisms is documented in individual lineages, a larger theoretical question remains unresolved: how do genome architecture, sequence constraints, and conserved pathways collectively shape evolutionary trajectories available for the grass family to adapt to diverse environmental stresses? Current knowledge is limited by sparse genome sampling and methodological constraints in comparative genomics. Hypotheses about global genomic features (e.g., adaptive GC shifts) have not been tested specifically in Poaceae. Likewise, functional studies of adaptive pathways and genes typically include several dozen species, focusing heavily on the ancient transition of Pooideae to cooler climates^13,38–42^. Consequently, the genomic architecture of many recent adaptive transitions among diverse grasses remains unexplored. Beyond limitations in sampling, inference of the functional consequences of sequence variation remains challenging because diverse classes of mutations (e.g., premature stops, frameshifts, splice disruptions, and missense substitutions) can converge on the same consequence through distinct molecular modifications, hindering unified inference in comparative genomic analyses^43–45^.

To address these gaps, we built on recent advances in genome sequencing initiatives^46–48^ and compiled an extensive collection of grass genomes, including 207 publicly available and 520 newly generated assemblies^49–53^. This dense phylogenetic sampling is expected to capture multiple independent environmental transitions at various times in the past, providing the natural replicates needed to test evolutionary associations at scale. Complementing this genomic resource, recent protein and nucleotide large language models (LLMs; e.g., PlantCAD- and ESM-series models) enable efficient and scalable sequence-based estimation of molecular features associated with gene function and evolutionary constraint^54–57^. By encoding sequences into high-dimensional embeddings that summarize latent biochemical and evolutionary properties, these models provide a practical framework for integrating diverse mutational effects in phylogenetic modeling.

Leveraging these genomic resources and protein and nucleotide LLMs, we test whether environmental adaptation across grasses is canalized through a limited number of mutational targets, such that independent colonization of similar environments repeatedly converges on the same genomic loci, stress-responsive pathways and genome properties.

## Results

### Repeated environmental transitions are widespread across the Poaceae phylogeny

To test whether extreme environmental niches have been occupied repeatedly by independent grass lineages, we studied the occurrence records and global environmental geodata for 561 focal Poaceae species. Assigning species to Köppen–Geiger climate zones^58–60^ based on their most frequently observed occurrences revealed that the sampled species grow across 17 distinct climate zones (Figure 1A). Most species occupy Tropical Savanna climate (Aw), but our sampling also spans semi-arid (BSh, BSk), arid (BWh), and multiple temperate zones (Cfa, Cfb, Csa), spanning the full spectrum of extreme temperature and precipitation regimes. We observed a wide range of growing environments across multiple subtribes, suggesting parallel adaptation among independent evolutionary lineages (Figure 1C).

**Figure 1.**
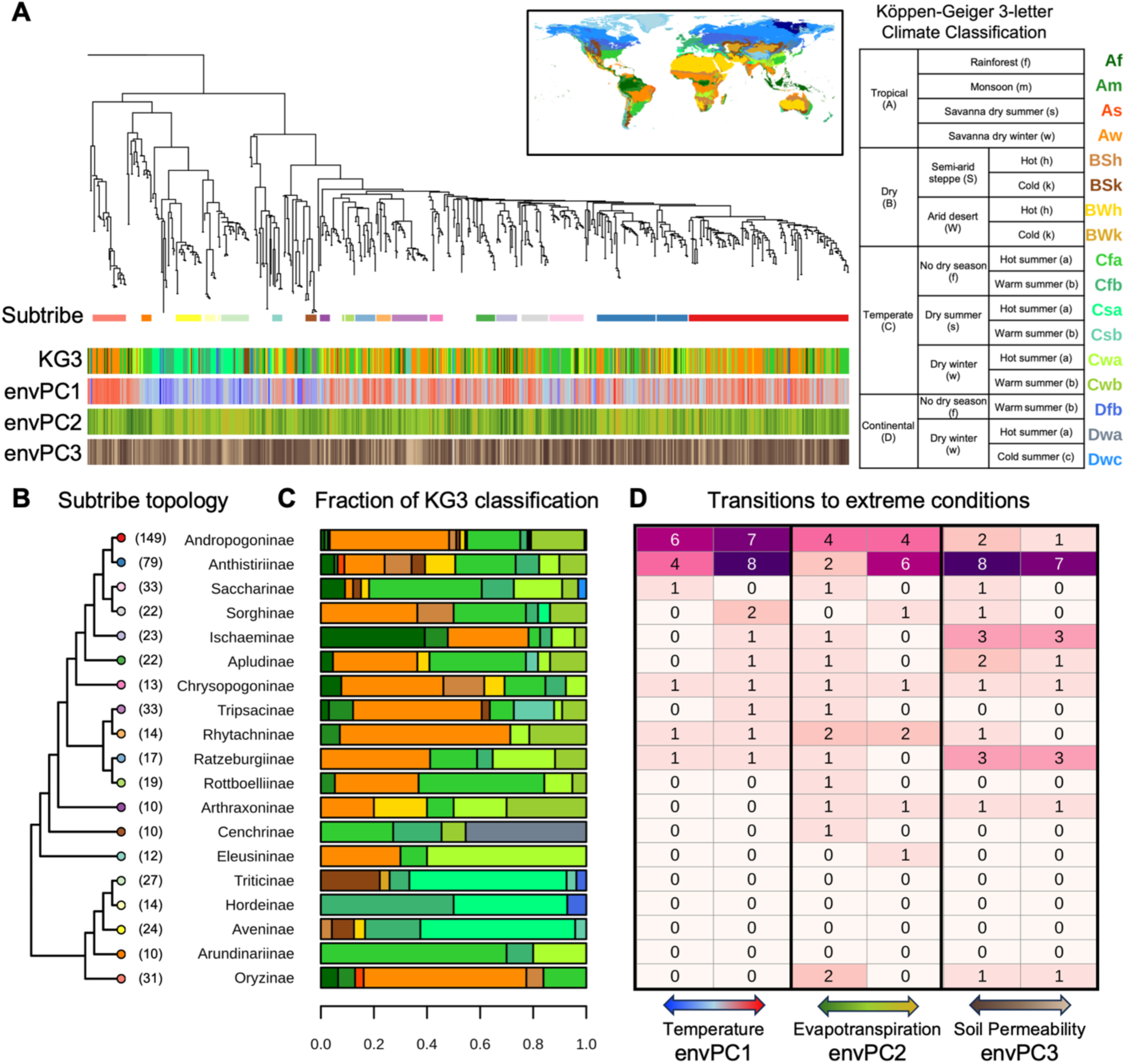
**Wide and repeated environmental transitions of grasses** A. The phylogeny of 687 Poaceae taxa. Tracks beneath the tree show, from top: major subtribe assignment (n ≥ 10; colors keyed to the subtribe labels in panel B); Köppen–Geiger climate classification of each taxon’s predominant habitat (KG3; colors keyed in the table at right); and scores on the first three environmental principal components (envPC1-3; colors keyed at the bottom of panel D): envPC1, temperate (blue) to tropical (red); envPC2, low (green) to high (gold) evapotranspiration; envPC3, high (dark brown) to low (light) soil permeability. B. The topology of major subtribes (n ≥ 10). The number in the bracket indicates sample size in each subtribe. C. The fraction of Köppen-Geiger habitat classification of the members in each major subtribe. D. The numbers of independent transitions toward the lower and upper tails (30%) of the envPC distributions.

To further quantify the habitat environmental variation among the studied species, we compiled 282 habitat-associated environmental variables and summarized them into principal components (envPCs; see Method details and Supplementary Figure S1). These envPCs represent orthogonal axes of habitat environmental variation across species (Supplementary Figure S1): envPC1 (27.0% variance) captures a tropical-temperate axis dominated by temperature and precipitation contrasts. envPC2 (17.8%) reflects a water-availability axis corresponding to an evapotranspiration gradient from tropical rainforest through monsoon, savanna, steppe, and desert habitats. envPC3 captures variation in soil permeability and thus the tolerance to waterlogging or nutrient loss. All three envPCs show significant macroevolutionary covariation with phylogeny (82.4%-93.1% explained variance), yet substantial variation persists within most subtribes (Figure 1A; Supplementary Figure S1). This pattern indicates that closely related species have diversified into contrasting environmental niches multiple times. Consistent with this, ancestral-state reconstruction of envPCs identified 17-30 independent transitions toward the lower and upper tails (30%) of the envPC distributions (Figure 1D).

Together, these analyses demonstrate that contrasting environmental niches have been colonized repeatedly across the grass phylogeny, providing natural replicates for dissecting the genomic basis of environmental transitions. The statistical power to detect convergent genomic signals depends critically on their convergence level and the magnitude of the underlying effects^61–63^.

## Dense phylogenetic sampling enables detection of convergent association signals

To evaluate whether our phylogenetic sampling provides sufficient power to detect convergent genomic signals in phylogenetic mixed models, we performed simulations with the empirical grass phylogeny and the transition frequencies inferred above (Figure 2 and Supplementary Table 1). In line with theoretical predictions^61–63^, the detection power scaled positively with the magnitude of the co-shifts and the frequency of co-shifting across lineages. Under the empirical transition scenario, our current sampling enables high power (>80%) to detect large and convergent signals while we are likely to miss some associations of moderate to low effects and low convergence. Notably, doubling the number of independent transitions captured with the same total sample size would increase the detection power of such signals by approximately 1.5-fold, underscoring the value of continued sampling across underrepresented grass lineages (Supplementary Figure S2).

**Figure 2.**
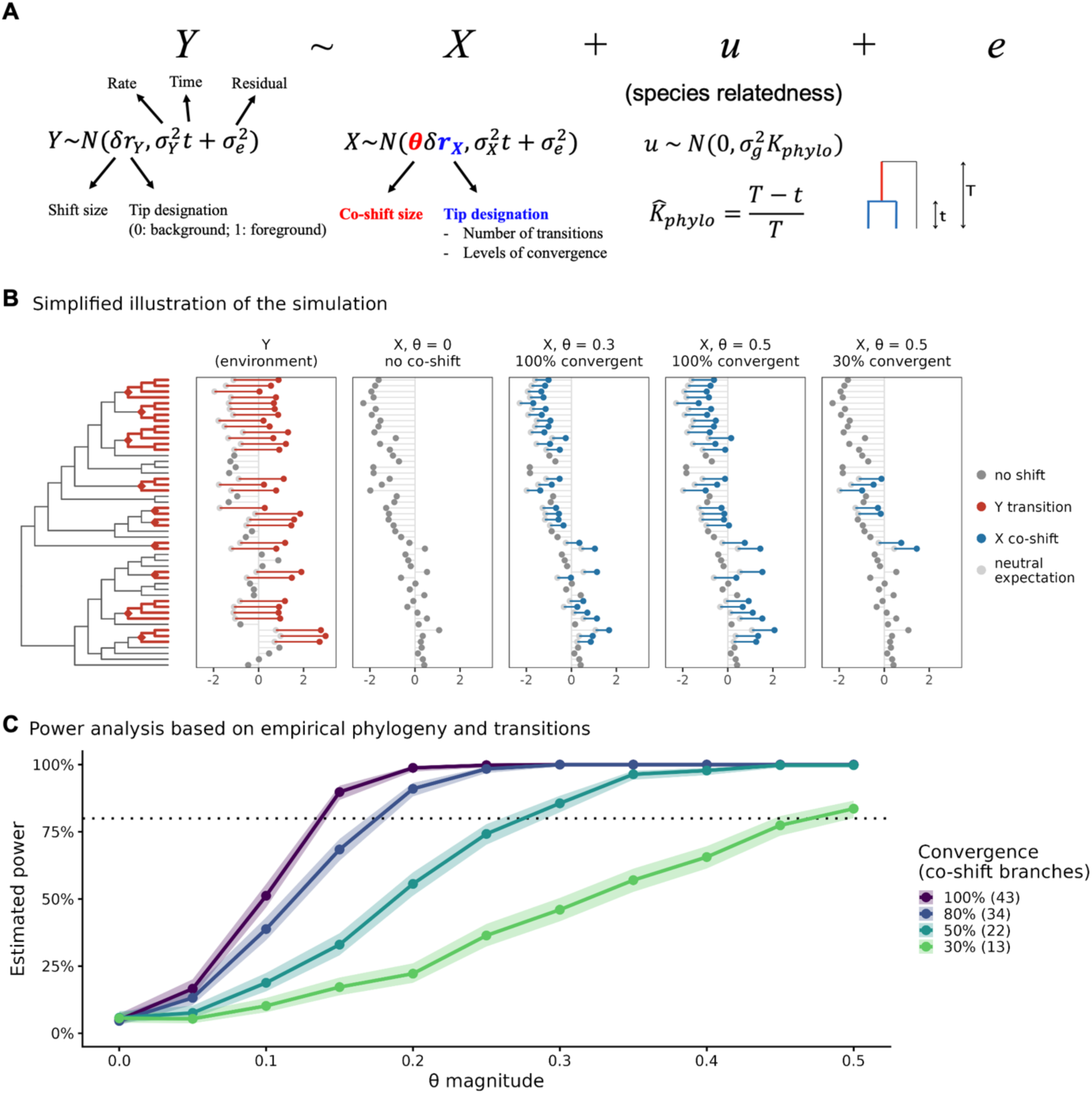
**Power analysis on phylogenetic mixed model** A. We simulated phenotypic (Y) and genotypic variables (X) that follow independent Brownian motion (BM) but share mean shifts at the transition branches in the phylogeny and evaluated the power of the phylogenetic mixed model in detecting such co-shift between the variables. B. Illustrative examples of the simulation. We showed a simplified phylogenetic tree of 50 tips and 10 transition nodes. Branches descendent from the transition nodes are labeled red. Besides the tree, simulated Y and X under different scenarios were shown. Grey dots indicate neutral values under BM while red/blue dots and lines indicate deviance from neutrality due to the simulated evolution shifts. We plotted scenarios where X showed (1) no co-shifts, (2) moderate co-shifts with 100% convergence, (3) strong co-shifts with 100% and (4) strong co-shifts with 30% convergence. These illustrate the effects of the two factors we manipulated in the simulations: relative co-shift size and convergence level (i.e., the frequency of co-shift across all transitions). C. The power curves under different simulated parameter configurations on the 43 empirical transition nodes to both envPC1 extremes. We tested varying relative co-shift sizes (x-axis: *θ*), and levels of convergence (different colors). The solid dots indicate average power across 500 simulations and shaded area denotes 95% confidence interval. The power of the phylogenetic mixed model increases as the convergence level and relative co-shift size increase.

### Repeated environmental shifts are accompanied by parallel modification of genomic properties

With the statistical properties of our analytical framework established, we first asked whether repeated colonization of similar environments was accompanied by convergent shifts in genome-wide molecular composition. Among the diverse grass species, the amino acid composition of orthologous protein-coding genes varies substantially (Supplementary Figure S3), implicating selection on the physiochemical properties of the encoded proteins. Indeed, quantification of key proteomic features revealed variation in nitrogen-to-carbon ratio, biosynthetic energetic cost, hydropathy, molecular weight, and density of the proteome-composing amino acids (Figure 3A). In parallel, GC content of coding sequences (CDS) ranged from 51.5% to 58.8% across genomes (mean = 54.8%; CV = 1.6%; Figure 3A). Given the central roles of transcription and translation^64^, these proteomic and nucleotide features are likely targets of selection imposed by distinct environmental stresses. Consistent with this, these traits exhibit heterogeneous macroevolutionary covariation with phylogeny (Figure 3B), suggesting differential selective constraints.

**Figure 3.**
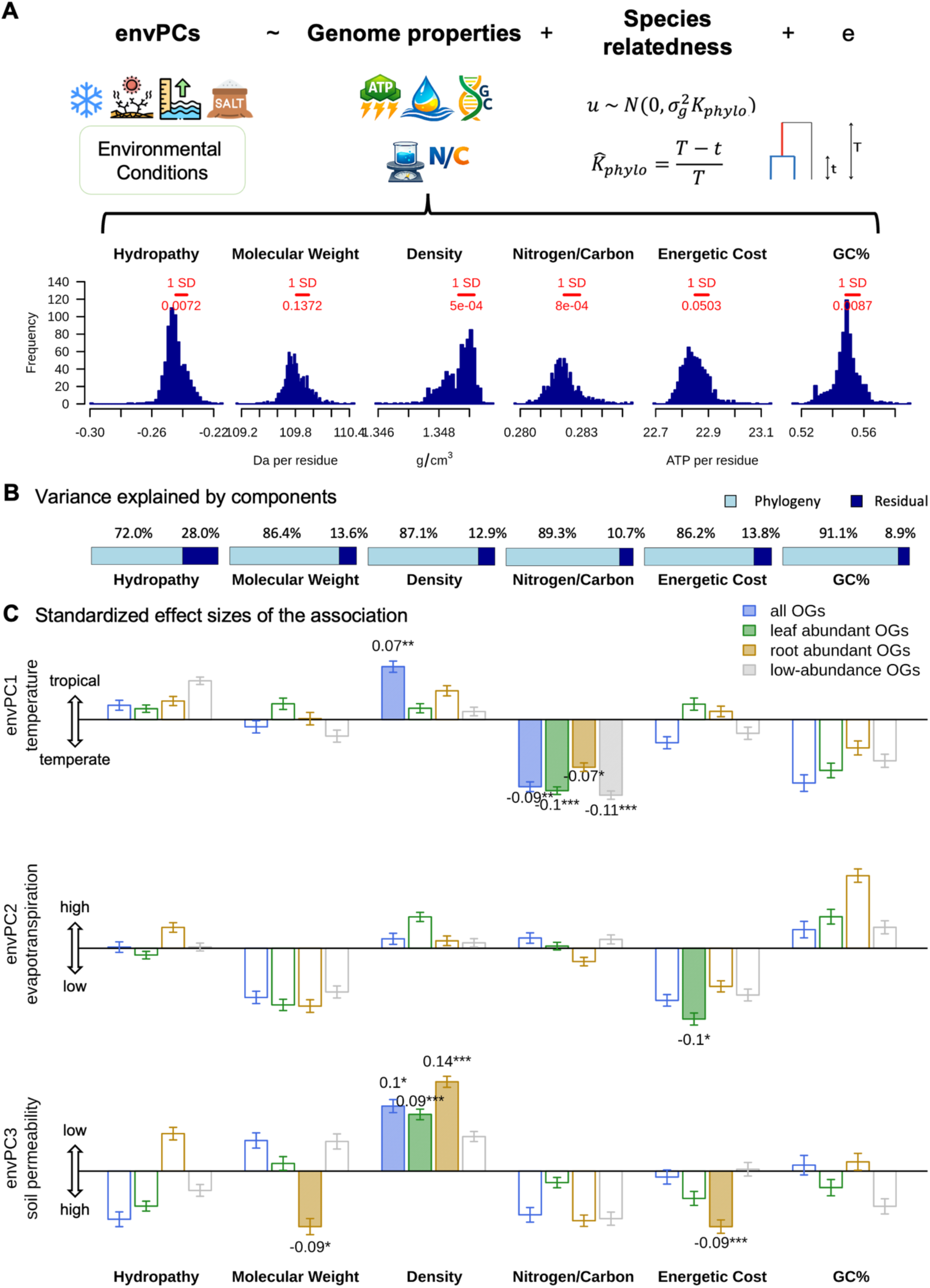
**Repeated environmental shifts are accompanied by parallel modification of genomic properties** A. The phylogenetic mixed model used to test the hypothesis that environmental variation is associated with the variation in the molecular and physiochemical features of grass genomes. The distributions of the studied molecular and physiochemical features are shown. One standard deviation of each distribution is indicated in red. B. The variance decomposition between phylogeny and residual. Null expectation of neutrally evolving traits would be fully explained by phylogeny (100% variance explained). C. Bar plots of the standardized effect size of each feature on each envPC (i.e., the magnitude of envPC shift in standard deviation by shifting one standard deviation of the feature). The same test was performed using different sets of orthogroups (OGs), including all OGs, leaf-abundant OGs, root-abundant OGs and low abundance OGs as indicated by different colors. Statistically significant effects are indicated by solid bars, and the effect sizes are labeled above or below the bars. The error bars were determined by 1,000 runs of phylogenetic permutation. The significance of the effect was determined by Wald test and confirmed with phylogenetic permutation.

To test whether environmental selection is associated with variation in these molecular and physiochemical features, we modeled each environmental axis (envPC1-3) as a function of individual molecular traits using phylogenetic mixed models. Relative to phylogenetically permuted null models^65^, multiple molecular traits showed significant associations with the first three envPCs (Figure 3C). Significant negative association was found between envPC1 and the amino-acid nitrogen-carbon ratio, indicating reduced use of nitrogen-rich amino acids in Tropical species. This is consistent with the stronger nutrient competition in warmer ecosystems^66–68^. As for the biosynthetic energetic cost, it is negatively correlated with envPC2 and envPC3 for leaves- and root-abundant proteins, respectively: Species experiencing severe evapotranspiration stress encoded energetically cheaper proteins in leaves, plausibly reflecting rapid tissue turnover under water stress^69,70^. Likewise, species inhabiting poorly drained soils evolved root-abundant proteins with lower biosynthetic energetic cost.

Together, these patterns are consistent with that environmental selection has repeatedly shaped core molecular features of grass genomes, driving convergent biochemical shifts across independent lineages.

### Nucleotide and protein large language models predict gene activity evolution de novo

Beyond genome-scale molecular and physicochemical features, environmental selection also operates at the level of individual loci. We therefore inferred and examined gene activity evolution across 19,613 orthogroups (OGs; see method details) in grass genomes. We define gene activity evolution as the change, across species, in the function of an orthologous gene as inferred from its sequence: spanning complete loss of the gene, truncation by premature stop codons, and accumulated missense substitutions that lead to functional modification. Consistent with the well-conserved synteny and gene content reported for Poaceae^71^, we observed (1) minimal gene loss (Supplementary Figure S4A); (2) rare occurrence of premature stop codons (Supplementary Figure S4B: only ∼5% of sequences per OG on average contains a premature stop codons); (3) pervasive purifying selection across orthologous genes (Supplementary Figure S4C: <5% of OGs exhibited a dN/dS ratio greater than 0.5 relative to the reference species, *Pharus latifolius*). The small subset of genes with elevated absence frequencies, accumulation of premature stop codons and molecular evolution rates is primarily associated with defense responses and pollen compatibility (Supplementary Figure S4A-C), traits often shaped by Red Queen dynamics^72–74^. Within each OG, homologous sequences from diverse grass species exhibited 10-150% variation in selective constraint, suggesting heterogeneous selective pressures and lineage-specific functional divergence (Supplementary Figure S4D). OGs with the most extreme divergence were enriched for functions related to post-transcriptional and translational regulation, highlighting the roles of these processes in evolutionary innovation.

In addition to comparative metrics based on multiple sequence alignments (MSAs), we applied nucleotide and protein large language models (LLMs) and evaluated a scoring framework that predicts gene activity de novo solely from sequence (Figure 4A). Using well-curated mutant datasets in Arabidopsis^44^ and Maize^75^, we found that our protein and nucleotide LLM scoring approach detects mutational effects as subtle as single-amino acid substitutions without requiring MSAs (Figure 4B). Mutations with experimentally validated phenotypic consequences produced significant reductions in model scores relative to wild type, whereas neutral mutations produced minimal changes (Figure 4C). Across all 19,613 grass OGs in this study, ESM2 scores effectively identified high-impact loss-of-function mutations, such as premature stop codons and frameshifts (Figure 4C and D), while PlantCAD was sensitive to elevated rates of missense substitutions (Figure 4C and E). These results demonstrate that LLM-derived predictors offer a scalable and sensitive proxy for gene activity evolution across hundreds of grass genomes.

**Figure 4.**
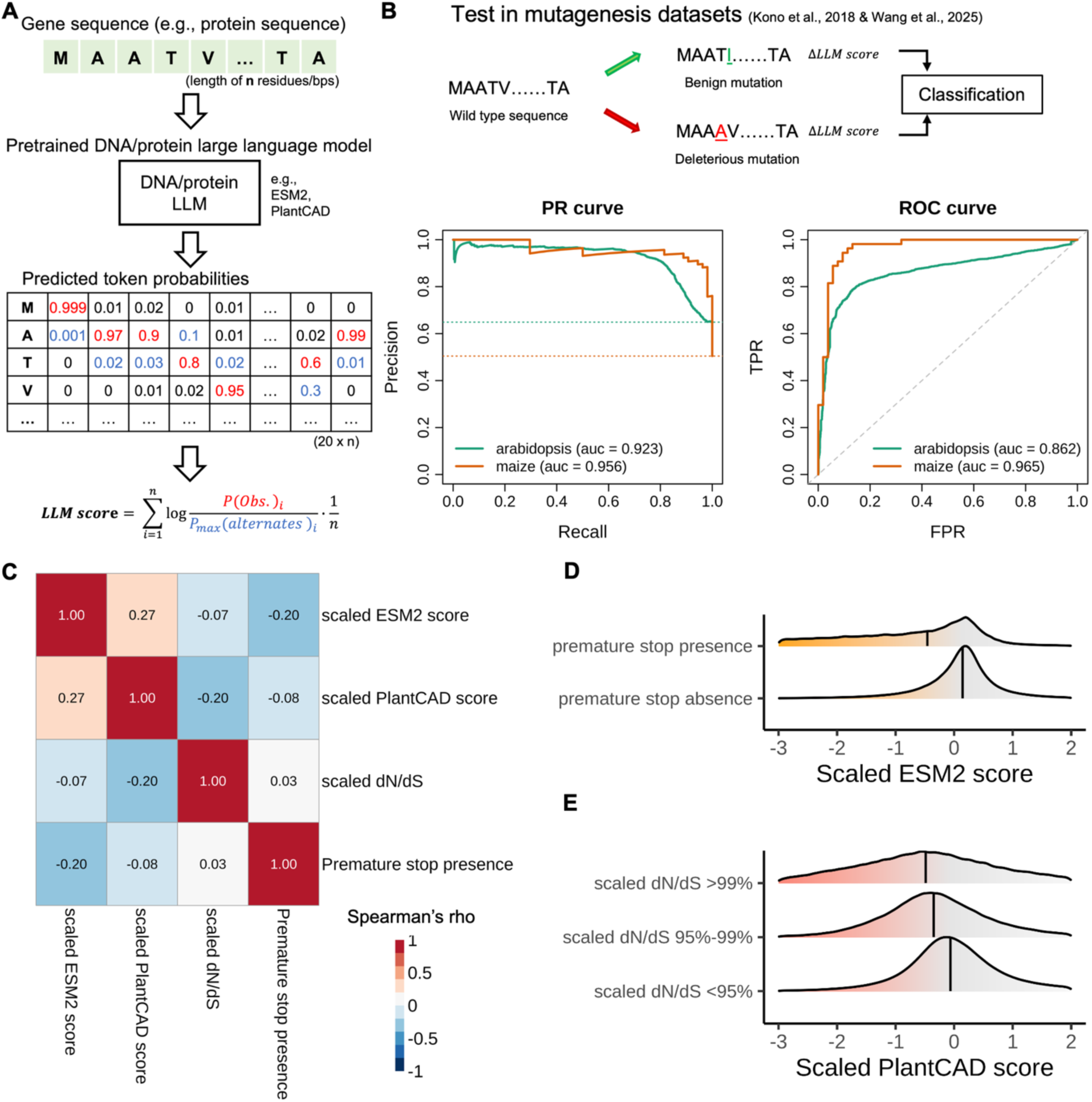
**Nucleotide and protein large language model (LLM) scores predict gene activity changes.** A. The derivation of LLM scores from zero-shot logit of nucleotide and protein LLMs. Based on the per-site prediction probabilities of all possible nucleotides/residues given the observed gene sequence context, we defined the LLM scores as: 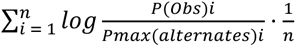 where i indicates the i^th^ residue/base pair in the sequence with a length of n. The lower the score is the less confident the model is in making the prediction, which implies lower functionality of the input sequence. B. Validation of the LLM scores using well-curated mutant datasets in Arabidopsis^44^ and Maize^75^. Precision-Recall (PR) and Receiver Operating Characteristic (ROC) curves are shown. C. Comparison between the LLM scores against MSA-based metrics of molecular evolution. Spearman’s rank correlation coefficients (rho) are shown. D. ESM2 scores effectively identified high-impact loss-of-function mutations, such as premature stop codons. E. PlantCAD was sensitive to elevated dN/dS (defined as above the 95% and 99% of the scaled dN/dS).

### Convergent selection responses of key genes are associated with repeated environmental shifts

Finally, we tested whether specific genes exhibit parallel activity shifts associated with environmental shifts. Using the phylogenetic mixed-model framework described above, we modeled each principal environmental axis (envPC1-3) as a function of gene-activity for each OG. Recognizing that the three MSA-based comparative metrics (presence-absence, premature stop codon occurrence and tip-to-reference dN/dS) and the two LLM-derived scores (ESM2 and PlantCAD) capture complementary aspects of gene activity, we fitted joint models including all predictors and compared full and reduced models using likelihood ratio tests (Figure 5A). Model significance was assessed using a phylogenetic simulation-corrected permutation test^65^.

**Figure 5.**
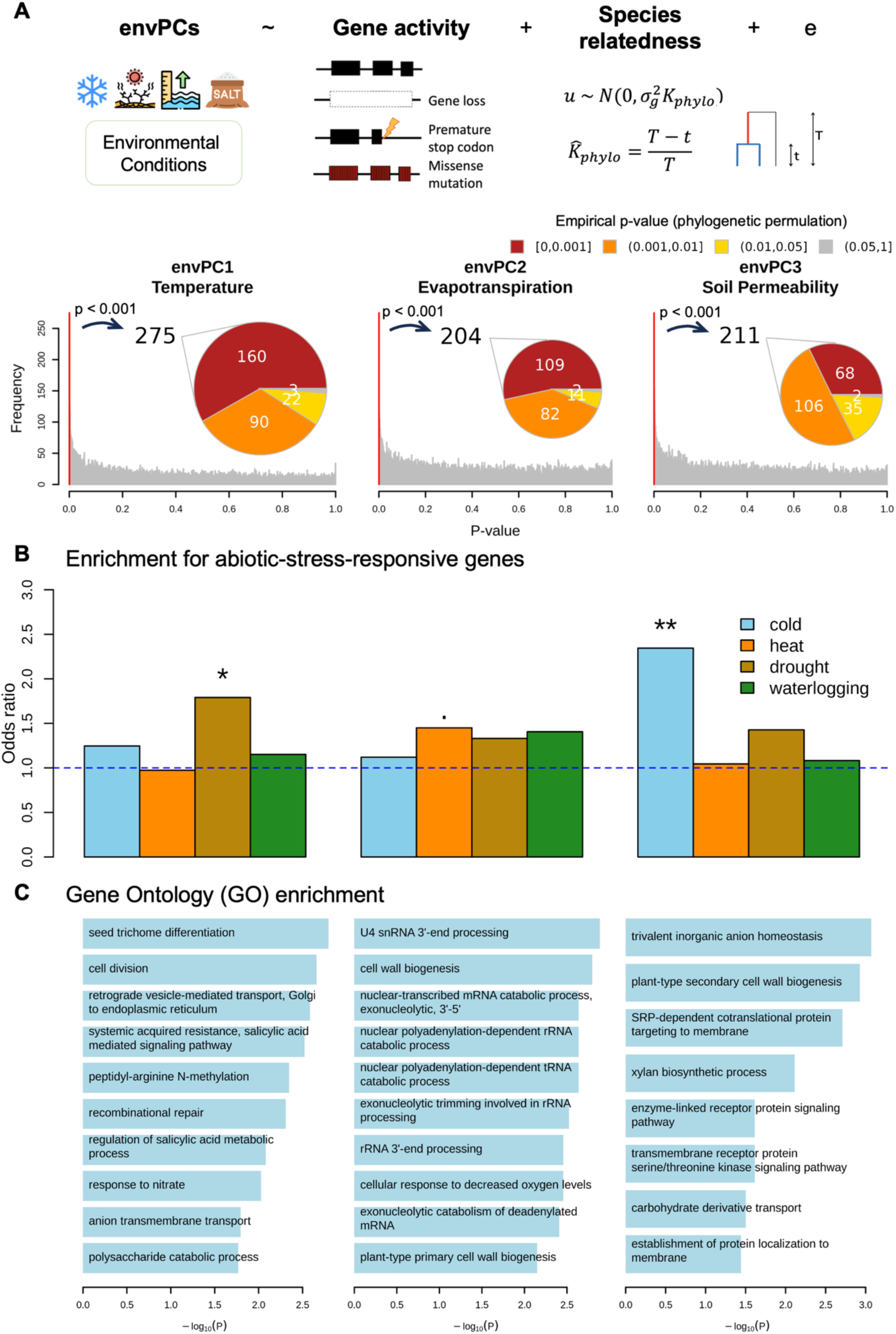
**Phylogenetic mixed model identified OGs convergently associated with top envPCs.** A. The phylogenetic mixed model used to test the hypothesis that environmental variation is associated with gene activity variation. The histograms of the asymptotic p-values are shown. On top of the histogram, the piecharts indicate the empirical p-values of the top candidate OG (asymptotic p-value ≤ 0.001) from phylogenetic permulation tests. Only the OGs with both asymptotic and empirical p-values ≤ 0.001 are considered significant. B. Enrichment of the candidate OGs against *a priori* abiotic-stress-responses genes. · p < 0.1; * p < 0.05; ** p <0.01. C. Gene ontology (GO) enrichment of the candidate OGs. Top 10 significant Biological Process (BP) terms ordered by significance are shown. Across panels, the three columns track the results for envPC1-3 respectively.

This analysis identified 160, 109, and 68 OGs significantly associated with envPC1-3, respectively (Figure 5A; asymptotic and empirical p-values ≤ 0.001; Supplementary Table 2). These associations were robust to phylogenetic sampling. Rankings of association strength (-log10P) were largely preserved when analyses were restricted to one accession per species (ρ = 0.79-0.91; Supplementary Figure S5), and when each of the ten subtribes with more than 20 taxa was removed in turn (envPC1; ρ = 0.78-0.99; Supplementary Figure S6). Candidate status was not associated with maximum protein abundance or with breadth of protein accumulation across tissues (Logistic regression: p = 0.71 and 0.70), indicating that detection was not biased toward highly or broadly expressed genes. Of the identified candidate OGs, 34.5-41.2% overlapped the curated set of *a priori* abiotic-stress-responsive genes across grasses (see Method details), with significant enrichment for specific stress categories (Figure 5B). Furthermore, they were enriched for biological processes implicating environmental responses (Figure 5C; Supplementary Table 3). For example, envPC1-associated OGs were enriched for salicylic acid signaling and cell division, highlighting their importance along the temperate-tropical axis. OGs associated with envPC2 implicated tRNA/rRNA processing, hypoxia response and cell wall restructuring under differential evapotranspiration stresses, whereas envPC3 highlighted membrane targeting and cell wall remodeling processes relevant to flood/nutrient leaching tolerance. We also detected significant enrichment for extracellular, chloroplast and mitochondria-localized proteins among the candidates (Supplementary Table 3), implicating preferential occurrence of environmentally associated modifications in specific subcellular compartments. These enrichments support the biological relevance of this screen for putatively adaptive genes across hundreds of genomes.

For a closer investigation, we applied two additional filters to identify a high-confidence subset of environmentally associated OGs. First, we retained only envPC-associated OGs that also exhibited significant parallel shifts in selection strength on transitional branches, as detected by RELAX, a codon-substitution method for testing shifts in selection intensity across predefined lineages^76^. Second, we intersected this set with a curated list of *a priori* abiotic-stress-responsive grass genes (see Method details). This filtering yielded a union of 27 OGs supported by concordant evidence from independent tests (Figure 6). Several of these candidates have previously been functionally characterized in abiotic stress responses in model systems. For example, orthologs of α-expansin proteins and RING-type E3 ubiquitin ligases showed consistent activity shifts between temperate and tropical biomes (envPC1). Overexpression of *ZmEXPA5* improves yield potential under drought conditions in maize, and its Arabidopsis ortholog, *AtEXPA2*, when overexpressed, also enhances tolerance to osmotic and salt stresses^77,78^. The rice ortholog of the RING-type E3 ubiquitin ligase, *OsDCA1*, acts as a coactivator of *OsDST* and confers drought and salt tolerance when knocked out^79^. Orthologs of xyloglucan endotransglucosylase/hydrolase (XTH) proteins were highlighted along the evapotranspiration gradient (envPC2), consistent with reports linking these enzymes to flooding responses in maize^80^. The subset also includes candidates with no described role in stress tolerance, such as a phosphoglycerate mutase (PGAM) ortholog whose predicted chloroplast-targeting signal shows significant variation associated with envPC1 (Supplementary Figure S7). Another interesting candidate is the envPC2-associated ZmKW9 ortholog, which encodes a pentatricopeptide repeat protein responsible for C-to-U editing of the plastid *ndhB* transcript within the NDH complex^81^.

**Figure 6.**
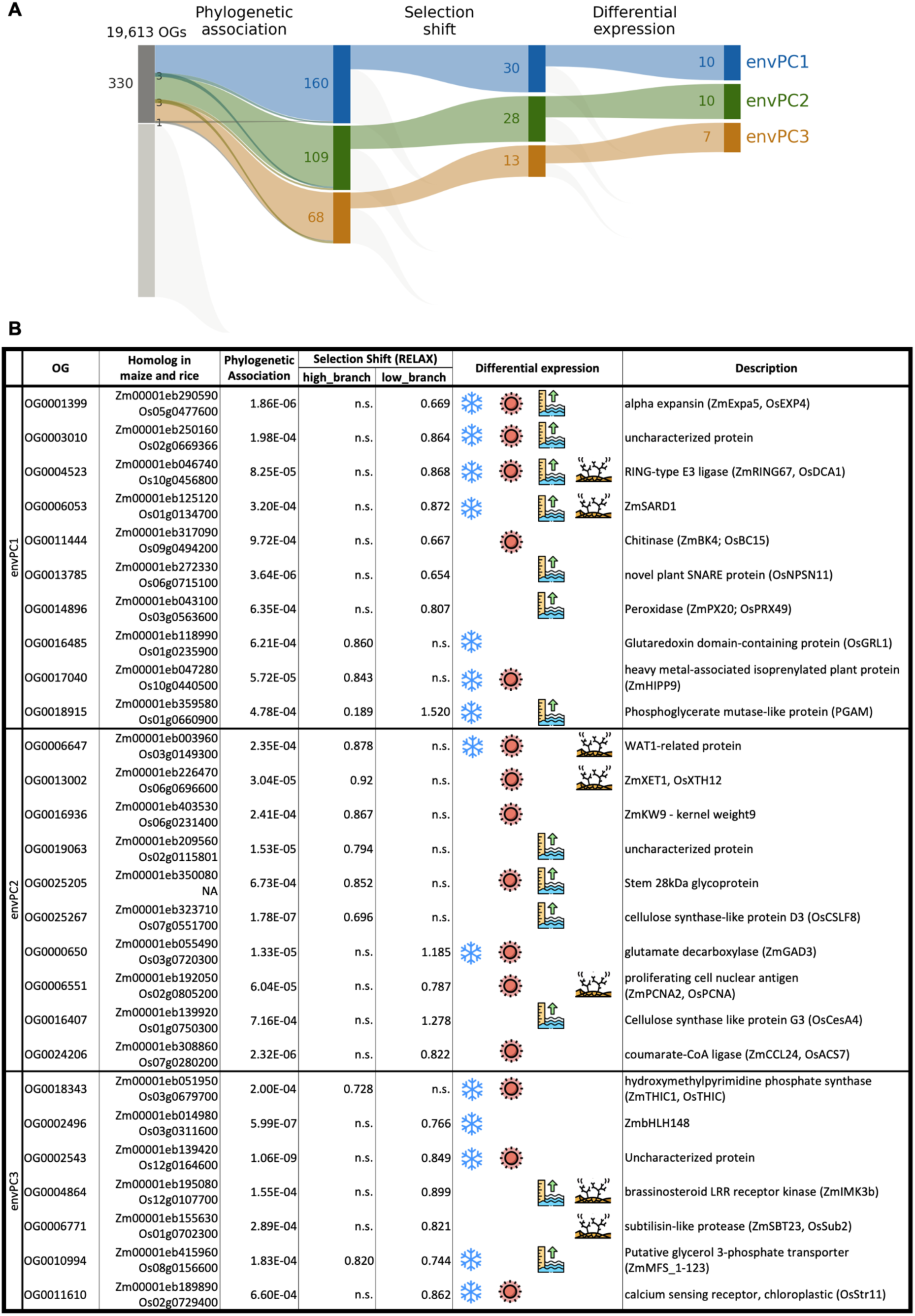
**High-confidence subset of environmentally associated Ogs** A. Three layers of independent tests to determine the 27 high confident environmentally associated OGs. The bar heights and ribbon widths use squared-root scale for visualization. A few OGs were detected in the association test for two envPCs. B. The 27 high confident associated OGs and the evidence of calling. For phylogenetic association, asymptotic p-values of the phylogenetic mixed model are shown. For the selection shift, k statistics of the significant RELAX tests (p-value < 0.01) are shown. k < 1 indicates relaxed selection while k > 1 indicates intensified selection. Symbols of environmental stressors are used to indicate the conditions in which each of the candidate OG changes their expression in ≥ 3 grass studies.

Beyond the high-confidence subset, the broader candidate set recapitulates the same functional themes. Consistent with the compartment enrichments above, 14 candidate OGs are predicted to be targeted to chloroplast membrane systems and 16 encode secreted or cell wall-localized enzymes, most without prior evidence of stress-responsive expression. Among the former are orthologs of PsbY, a subunit of photosystem II, and of CURVATURE THYLAKOID 1A (ZmCURT1A; Zm00001eb190110), which shapes thylakoid membrane architecture and is required for the onset of photosynthesis^82,83^. The set also includes an ortholog of ZmAGAL4, linked to chloroplast lipid metabolism under osmotic stress^84^. Together, these point to the organization and stability of the photosynthetic membrane as recurrent targets of environmentally associated changes. Among the cell wall localized enzymes, we identified another expansin ortholog (Expansin-B4; Zm00001eb000370), which co-localizes with a QTL for ear diameter in maize^85^, and an ortholog of AtLOTR1, a putative extracellular protease whose loss of function disrupts Casparian strip formation and compromises the apoplastic diffusion barrier in Arabidopsis roots^86,87^. These implicate cell wall extensibility and apoplastic barriers, the structures governing water and solute exchange between the plant and its environment, along the same environmental axes.

## Discussion

The mutational target size for selection responses fundamentally determines the repeatability of the evolutionary trajectories in independent lineages^88,89^. One or a few predictable routes of genetic changes would be taken when the mutational target size is small, as in the convergent loss of shattering genes during cereal domestication^90^ or the melanin pigmentation evolution in vertebrates^91^. When the mutational target size is large and genetic redundancy is pervasive, convergent targets of selection are rarely detectable even between replicates of an evolution experiment^92^. Analyzing hundreds of genomes from species occupying diverse climates, our results suggest that the ecological dominance of grasses reflects complex but constrained mutational targets as convergent association signals were detected while the signals span hundreds of loci, multiple pathways and genome-wide molecular composition.

The genome-scale signals tie the cellular metabolic processes with organismal adaptation. Because protein synthesis represents one of the largest energetic and nutrient investments in plant cells^64^, selection imposed by resource-limited environments would favor proteomes that are metabolically cheaper to synthesize and maintain^14,15,23^. Consistent with this prediction, species inhabiting environments with strong nutrient competition (as indicated by high envPC1 values), high evapotranspiration (high envPC2 values) or poor soil drainage (high envPC3 values) tended to encode proteins with reduced nitrogen demand and lower biosynthetic energetic cost (Figure 3). Notably, GC content of the coding nucleotides showed a comparatively weaker environmental association than the amino acid features, suggesting stronger selection on amino acid than nucleotide compositions. This is consistent with the substantially larger metabolic investment required for translation relative to transcription and DNA replication^64^.

At the gene level, candidate OGs were enriched for extracellular, mitochondrial and chloroplast proteins (Supplementary Table S3). Extracellular proteins mediate stress perception and cell wall remodeling^93^, and our candidate OGs include expansins and XTHs. Elevated expansin and XTH activity, which maintains wall extensibility, is among the most consistent cell wall responses to abiotic stress^94^, and XTHs are enriched among genes with parallel divergence between foothill and alpine ecotypes of *Arabidopsis arenosa*^95^. The enrichment of photosynthesis and mitochondrial electron transport genes among climate-associated loci was also documented in *Arabidopsis* populations^96^. Photosynthesis is temperature-sensitive^97^, and Rubisco kinetics show convergent adjustment to species’ thermal history^98^. Yet because plastid genes evolve slowly^99^, the nuclear-encoded components offer additional and potentially more accessible routes for tuning plastid function through complex assembly, RNA processing and protein import^100^. Our candidate OGs exemplify these routes: CURT1A organizes the thylakoid membrane; PsbY is a part of the photosystem II complex; KW9 edits plastid ndhB transcript, and PGAM carries chloroplast transit peptides whose predicted targeting probability varies with environment (Supplementary Figure S7). Notably, the NDH complex appears as a common target of adaptive changes: natural variation in the nuclear-encoded maize *ndhm1* alters photosynthetic efficiency and cold tolerance in European landraces^101^.

These targets offer flexible routes for change at limited cost: extracellular and membrane proteins evolve faster than intracellular, soluble ones^102,103^, most chloroplast proteins are encoded by genes transferred to the nucleus, which evolves faster than the plastome^99,100^; nuclear-encoded editing factors determine amino acids in plastid-encoded proteins and co-evolve rapidly with their target sites^104,105^, and transit peptides are modular N-terminus sequences that permit change in targeting while retaining enzymatic function^106,107^. Such flexibility could be the key feature to expand the range of accumulated variants available for selection as environments change, predisposing these genes to become recurrent targets of selection^108^.

Our ability to detect these phylogenetic association signals at scale rests on a methodological departure from classical “rate-shift” approaches that require branch-wise rate estimation (e.g.,^76,109^). By modeling environmental variables directly against sequence-inferred gene activity, we take the “state-state” association strategy^62,110^ that is both computationally scalable and statistically well-suited to the dense phylogenomic sampling available across Poaceae. Particularly, the use of nucleotide and protein large language models^55,56^ serves as an efficient supplement to the MSA-based comparative metrics in summarizing independent mutational events into a unified estimate of gene activity “state.” The statistical power of this approach depends critically on the number of independent evolutionary transitions sampled. Our current sampling captured 17-30 transitions per environmental axis, and simulations indicate that doubling this number would increase power by approximately 1.5-fold (Supplementary Figure S2). Expanding genome representation in currently underrepresented clades is therefore the clearest path to improved sensitivity. For example, Aristidoideae, Danthonioideae and Micrairoideae are not represented, and Panicoideae outside Andropogoneae are represented by only 18 species despite encompassing >10 origins of C4 photosynthesis as these lineages colonized open, often drier habitats^7,10^. Targeted sampling of these clades would add independent environmental transitions absent from the current analysis.

In conclusion, the core finding of this study is that environmental adaptation in grasses is canalized through a genetic architecture that, though complex, is convergent enough to be partially predictable. This has direct implications for how evolutionary genomics connects to practice. Because adaptation involves hundreds of loci and genome-wide molecular composition shifts, precision editing of single genes for environmental resilience may be of limited utility for breeding or conservation though disentangling drivers from passengers among the candidates may identify a few core targets worthy of editing. A more productive application of this predictive framework is upstream — guiding genomic prediction, assisted gene flow, and targeted recombination of natural diversity by identifying genomic configurations suited to particular environmental contexts. As grasses underpin both global food production and many of the world’s most threatened ecosystems, the ability to anticipate evolutionary responses to environmental change is no longer only a scientific question but a practical imperative for food security and conservation.

### Limitations of this study

Several limitations remain. First, current constraints on assembly contiguity and language model context length preclude detection of regulatory evolution, gene family dynamics, and polyploidy-driven dosage effects^111–119^ — mechanisms through which canonical stress-response families such as CBF/DREB operate primarily^31,32^. Our candidate OG set therefore represents a specific, if incomplete, layer of the genetic landscape, enriched for loci where coding sequence variation is the primary mechanism. Future development of low-cost long-read sequencing technologies and long-context genomic language models will likely lift these constraints, enabling a more comprehensive understanding of the genomic basis of Poaceae’s ecological dominance. Second, environmental niches were characterized from occurrence records, which carry a certain degree of geographic sampling bias^120^. Targeted flora investigation in under-studied regions (e.g., South and Southeast Asia) is needed to alleviate potential bias in information. Finally, none of the candidate genes in this study has been functionally tested. Such validation is challenging here as most sampled species are not amenable to genetic transformation. Furthermore, the layered and polygenic architecture we observe implies that individual alleles have small, background-dependent effects that may not produce measurable phenotypic changes when modified individually. A more tractable test would be whether prioritizing these loci improves genomic prediction of performance across environments in populations derived from crosses with environmentally divergent relatives.

## Resource Availability

### Lead contact

Further information and requests for resources and data should be directed to and will be fulfilled by the lead contact, Sheng-Kai Hsu

### Materials availability

Newly collected Andropogoneae plant vouchers associated with this study were deposited at Missouri Botanical Garden Herbarium; and for Australian wild collections, duplicates were deposited at Australia’s National Herbarium in Canberra. This study did not generate any new, unique reagents.

### Data and code availability

• The associated raw sequence data are available through sequence read archive (SRA) and the assemblies generated will be released through Ag Data Commons at https://figshare.com/s/eb35a1d65e4eb6be7c79. Derived occurrence datasets from BIEN and GBIF.org are available at https://zenodo.org/records/14968186.

• Source scripts, analytical notebooks and derived datasets used will be available at https://github.com/maize-genetics/p_phyloGWAS.

• Any additional information required to reanalyze the data reported in this paper is available from the lead contact upon request.

## Acknowledgements

Obtaining plant material for this project would not be possible without our many collaborators in the field and herbaria. We are grateful for their expertise, willingness to aid in field collecting, and sharing of precious herbarium material. In no particular order, we thank the following individuals, governmental organizations, and research institutions: *Individuals*: Taylor AuBuchon-Elder led the germplasm collection and plant maintenance with help from the following individuals: Donald Danforth Plant Science Center Plant Growth Facility staff, Rémy Pasquet, Cassiano Welker, Chrissy McAllister, Pat Minx, Bess Bookout, Maria Vorontsova, Jordan Teisher, Pete Lowry, Sarah Mathews, Richard Jobson, Tim Teetaert, Julie Pelc, Rev. Dennis Testerman, Russel Juelg, James Cole, Ron Day, Courtney Angelo, Chris Matson, Douglas Rogers, Michael McKain, Marshall Shaw, Arthur Stiles, Chuck Byrd, Kyle Dillard, Robert Findling, Nancy Sferra, Jonathan Bailey, Lynn Riedel, Brian Anacker, Kirsti Harms, Brandon Crawford, Charlotte Reemts, Matt McCaw, Kevin Thuesen, Michelle Bertelsen, Ryan Middleton, and Wesley Newman. Plant vouchers from available material were deposited at Missouri Botanical Garden Herbarium; and for Australian wild collections, duplicates were deposited at Australia’s National Herbarium in Canberra. *Herbaria* (silica dried leaf tissue and herbarium specimen tissue): Missouri Botanical Gardens, Royal Botanic Garden Kew, Muséum national d’Histoire naturelle, Australia National Herbarium, Queensland Herbarium, Northern Territory Herbarium, and National Herbarium of New South Wales. *Governmental organizations* (permits, permissions, and live plant or seed material): Australian National Government, New South Wales National Parks and Wildlife Service, Queensland Parks and Wildlife Service, Queensland Department of Environment Service, Northern Territory Government, Northern Territory Department of Natural Resources, Victoria Department of Environment, Land, Water, and Planning, North Carolina Department of Agriculture and Consumer Services, NC Plant Conservation Program, US National Park Service, US Department of Agriculture, City of Boulder Open Space and Mountain Parks, Florida Park Service and Department of Environmental Protection, and City of Austin Water Quality Protection Lands. *Non-profits, NGOs, research institutes* (permits, permissions, and live plant material): Katy Prairie Conservatory, Lady Bird Johnson Wildflower Center, The Nature Conservancy (Alabama, Arizona, Florida, Maine, Manitoba, Missouri, New Mexico, Texas), Native Prairie Association of Texas, University of Alabama, New Jersey Conservation Foundation, and University of Puerto Rico at Mayagüez. All collections were done in compliance with the Nagoya Protocol, with permits as required by local authorities. We acknowledge and honor the many custodians and stewards of wild and domesticated grass diversity worldwide who have shaped the study, preservation, and cultivation of grasses. We thank Jean-Luk Jenik for his advice on early implementation of mixed linear models in this study. Inari Agriculture provided financial support for the sequencing of six genomes, and Corteva Agriscience provided financial support for the sequencing of 10 genomes included in this study. This project is supported by the NSF PanAnd Grant Award #1822330 and USDA-ARS (8062-21000-043-00-D and 8062-21000-052-004-A).

Computational resources and data management were provided by the Bioinformatics Facility (RRID:SCR_021757) at the Cornell Institute of Biotechnology. Additional computations were performed on the Ceres and Atlas supercomputers at the SCINet HPC Consortium project and/or the AI Center of Excellence of the USDA-ARS, project numbers 0201-88888-003-000D and 0201-88888-002-000D. AJS is supported by NSF GRFP DGE-2139899. MCS is supported by NSF PRFB 1907343. EO is supported by the United States-Israel Binational Agricultural Research and Development Fund through the Vaadia-BARD Postdoctoral Fellowship Award No. FI-628-2022. BAK is supported by the Schlumberger Foundation Faculty for the Future program.

## Author contributions

S-KH, AJS, COH, MCR, and ESB conceived the research. TMA-E and EAK collected the materials. TMA-E, EAK, S-KH, AJS, COH, MCS and MCR curated the collected data. S-KH, LJB, EO, YL, BAK and JOO-R collected and curated expression datasets on abiotic stress responses. MCR managed the collected data. S-KH, AJS, and COH performed the formal analysis with domain support from GC-N, ZRM, MCS, TW, JZ, W-YL, HDD, QW, JOO-R and MP. S-KH, MCR and ESB managed the project. S-KH drafted the initial manuscript, and all other authors edited and approved the submitted manuscript.

## Declaration of generative AI and AI-assisted technologies

During the preparation of this work, icon design is facilitated by resources on Flaticon.com, and D-ALLE. We disclose the usage of OpenAI ChatGPT, Google Gemini and Anthropic Claude models to assist with code development and English editing. All AI-generated code and text was manually verified and edited, and the authors take full responsibility for the contents.

## STAR Methods

### Method details

#### Genome assemblies and quality check

We compiled a dataset of 727 genome assemblies representing 589 diverse grass species. Sampling was concentrated in the tribe Andropogoneae, for which we generated assemblies for as many taxa as available germplasm, field collections and herbarium specimens allowed. These include 33 newly generated long-read assemblies^53^ and 430 additional short-read assemblies released in this study using the high-throughput assembly pipeline described^52^. Outside Andropogoneae, we incorporated 207 publicly available genome assemblies from NCBI, Phytozome and CoGE, together with 57 assemblies generated from publicly available short-read data in the Sequence Read Archive^49^. Taxa were included based on the availability of material or data and were not selected according to their environmental niches. Up to five accessions were retained per species. Taxonomy, data source and sequencing technology for each assembly are listed in Supplementary Table 4. Obvious microorganismal contamination and misidentified assemblies were excluded as described^49^. We assessed assembly completeness using contig N50, median length of gene-bearing contigs, and a BUSCO-like assembly metric designed specifically for grasses^121^. Among these assemblies, 20 lack occurrence records (n < 6) and another 20 were flagged with anomalously high occurrence rates (>15%) of frame shift mutations and premature stop codons (See sections below for details). These 40 samples were excluded, leaving 687 assemblies representing 561 species in all subsequent analyses in this study.

#### Orthogroup construction

Across the 687 genome assemblies, 32 representative high quality long read assemblies were selected to construct orthogroups (OG). In order to avoid potential annotation biases, we ran Helixer^122^ to annotate each of the representative genomes and extracted the protein sequences. Based on protein sequence homology, OGs were constructed using OrthoFinder v2.5.4^123^. In total, we obtained 22,363 OGs with homologous sequences represented by more than 8 of the representative genomes. From the multiple sequence alignment of each OG, we reconstructed the ancestral protein sequence using the R/phangorn package (v2.12.1)^124^. The ancestral sequences of the OGs (Supplementary Table 5) were used to query the orthologous sequences in all 687 genomes by miniProt v0.13.0^125^ to build the full OGs. Since the assemblies were analyzed as deposited or assembled, without phasing into subgenomes or haplotypes, each ancestral protein was aligned to the whole assembly with miniProt, and every alignment passing miniProt’s output thresholds was retained as a separate copy with its own coding and protein sequence. We did not attempt to classify additional copies as homeologs, alleles or paralogs because phased haplotypes are unavailable for most assemblies. Since OG membership is defined by similarity to a reconstructed ancestral protein, homeologs, recent paralogs and alleles (if assembled) of a gene fall into the same OG by construction rather than forming spurious OGs. We used MAFFT v7.520^126^ to align the nucleotide and amino acid sequences for each OG. This pipeline is illustrated in Supplementary Figure S8. Maize proteome data^127^ were used to define tissue-abundance of the OGs.

#### Phylogenetic inference

Phylogenetic relatedness among the 687 studied genomes was constructed based on the angiosperm 353 loci^128^. We identified the orthogroups homologous to the angiosperm 353 loci using miniProt v0.13.0^125^ and generated gene trees for each of the orthogroups using RAxML v8.2.12 (GAMMA + GTR)^129^. ASTRAL-Pro v2^130^ was used to reconcile the species tree based on the gene trees. We used *Pharus latifolius* (Pharoideae), an early-diverging grass lineage outside the BOP–PACMAD clade that contains all focal taxa, as the reference for rooting and comparative analyses. This phylogenetic tree is available as Supplementary Figure S9. We generated a matrix of relative shared branch length among any pair of tips in the species tree to represent the relatedness between them^110^. This matrix (hereafter Kphylo matrix) is included in our phylogenetic mixed model to account for shared macro-evolutionary history among species.

#### Habitat environmental characterization

We leveraged an environmental characterization pipeline previously described^50^ to characterize the environmental niches of the studied species. For each species, we retrieved the geographical coordinates of occurrence from BIEN^131^ and GBIF.org^132^ and obtained various environmental features in which the species occurs. As mentioned, 20 assemblies without habitat environmental characteristics were excluded from subsequent analysis. The variation in habitat environments amongst the diverse species was summarized into environmental principal components (envPCs) and the top three envPCs were modeled as response variables in the phylogenetic mixed model. Additionally, we modeled the ancestral states of each envPC using R/ape (v5.8-1) and identified independent transitions to extreme conditions (defined here as the lower or upper 30% of the envPC distributions).

#### Amino acid composition and GC content

Across plant species, adaptation to divergent ecological niches is hypothesized to involve genome-wide molecular modifications to accommodate resource availability and competition. Particularly, proteins and the encoding DNA/mRNA are the most prominent selection targets given their central roles in biology. Hence, for each taxon, we calculated the amino acid composition and the derived physiochemical features, including molecular weight, density, hydropathy, Nitrogen-Carbon ratio and biosynthetic energetic cost, of all orthologous protein sequences using R/canprot package (v2.0.0). In addition, GC content was calculated using SeqKit (v0.15.0)^133^ for all coding sequences of the orthologous genes in each taxon.

#### Gene activity prediction of orthogroups

As species diverged and adapted to varying environmental niches, selection constraints differentiated, allowing functional divergence among orthologous sequences. When a gene confers no fitness benefit in a certain environment, relaxation of selection could lead to the excess accumulation of nonsynonymous mutations, occurrence of premature stop codons and ultimately complete loss of the gene, leaving sequence signals of changes in its activity. We focused our analysis on 19,613 testable orthogroups; these were orthogroups that were: (1) present in our reference species, *Pharus latifolius*; (2) present in at least 100 genomes in our focal clade. Based on the multiple sequence alignments of these orthogroups, we assessed the presence and absence of sequences for each taxon and identified premature stop codons that truncate the protein product by 10% in length. Unusually high occurrence rates (>15%) of frame shift mutations and premature stop codons were observed in 20 assemblies. We attributed this to sequencing and assembly errors and thus excluded these 20 assemblies, leaving a total of 687 genomes in subsequent analyses. Furthermore, we calculated dN/dS ratio for each sequence compared to *Pharus latifolius*, using R/MSA2dist (v1.2.0). Very few sequences with this “tip-to-reference” dN/dS ratio > 5 were attributed to alignment errors and treated as absent in subsequent analyses. We considered this “tip-to-reference” dN/dS estimation as a simplified implementation of the use of “root-to-tip” dN/dS in previous phylogenetic comparative studies (e.g.,^134^). In addition to the multiple sequence alignment (MSA)-based estimates, we also integrated nucleotide and protein LLMs, PlantCAD (kuleshov-group/PlantCaduceus_l32)^55^ and ESM2 (facebook/esm2_t33_650M_UR50D)^56^ to assess the function of orthologous gene sequences. These biological sequence LLMs predict nucleotide base pairs and/or amino acid residues with high confidence in the context of functionally/evolutionary conserved sequences. The average ability of the model to make unambiguous predictions for all sites in each sequence would reflect the functional constraint of this sequence. Hence, we defined a LLMs score based on the per-site prediction probabilities of all possible nucleotides/residues given the observed gene sequence context as following:

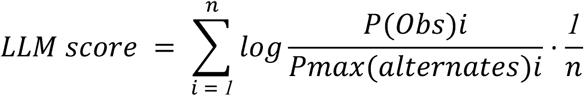

Where i indicates the ith residue/base pair in the sequence with a length of n. The lower the score, the less confident the model is in making the prediction, which implies lower functionality of the input sequence. Presence-and-absence was scored as the presence of any copy in an assembly and is therefore unaffected by copy number. Premature stop codon occurrence, tip-to-reference dN/dS, ESM2 and PlantCAD scores were computed for each copy separately.

#### Validation of LLM score

To validate the efficacy of the LLM scores in pinpointing loss in gene activity, we tested whether the proposed approach is able to distinguish single-amino acid substitutions with and without experimentally validated phenotypes in well-curated mutagenesis datasets^44,75^. For each gene, we calculated the LLM scores for all pairs of wild type and mutated sequences with a single-amino acid substitutions and derived the difference (*ΔLLM score*). Then, we evaluated whether the *ΔLLM score*s distinguish the mutation classes using the Precision-Recall (PR) and Receiver Operating Characteristic (ROC) curves and the area under the curves.

#### Power analysis on phylogenetic mixed model

Using the ASREML-R package (v4.2.0)^135^, we performed phylogenetic mixed modeling analyses^61,62,110,136^ to test the hypotheses in this study. ASREML-R implementation of the linear model analysis yields nearly identical results as the lambda model implemented in R/phylolm package (v2.6.5)^137^ for both estimated parameters and model significance (Spearman’s ρ > 0.998; Supplementary Figure S10).

Prior to hypothesis testing, we performed a simulation to assess the power of this approach. Given the empirical grass phylogeny and shift nodes in this study, we simulated a trait (Y) following Brownian motion with a fixed rate 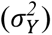 and mean shift (*δ*) below the transition nodes (as indicated by *r_i_*). In addition, a random error 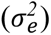 is introduced.

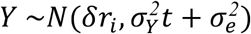

Where t is time; 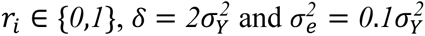

Similarly, we simulated genetic variables (X) under independent Brownian motion with a fixed rate 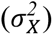 that exhibit smaller mean shift along the same transition branches (scaled by *θ*)

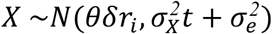

Where t is time; 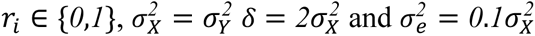

We simulated four transition scenarios: (1) empirical transition for envPC1 as inferred in this study, random transition of (2) the same, (3) twice or (4) half the empirical ones. Furthermore, we simulated various co-varying effect sizes (*θ*) from 0 to 0.5 with a step of 0.05 and different levels of convergent evolution by subsampling the transition branches to 100%, 80%, 50% or 30% of all transitions.

For each parameter configuration, we evaluated the frequency to detect the covariation between Y and X across 500 simulations as the power for the model below:

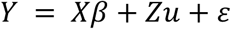

Where Y and X are the simulated phenotypic and genetic variables as described. *β* is the estimated effect size. *u* stands for the random effect term for phylogenetic relatedness and follows 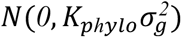 is the indicator matrix for *u*. *ε* is the residual.

#### Hypothesis testing using phylogenetic mixed model

Two hypotheses were tested in this study: (1) Environmental transition is associated with predictable, convergent modifications in genome-scale molecular/physiochemical properties; (2) Environmental transition repeatedly involves specific orthologous genes.

For the first hypothesis on environmental transition and genome-wide molecular modification, we modeled the top three envPCs against different physiochemical properties of the proteinogenic amino acid and coding DNA separately as follows:

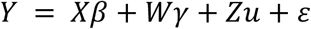

Where *Y* is the envPC, *X* is the focal physiochemical property as one of the following: molecular weight, density, hydropathy, Nitrogen-Carbon ratio, biosynthetic energetic cost and GC content. *W* stands for the nuisance factors including sequencing technology and assembly completeness + duplication level. *β* and *γ* are the fixed effects of predictor term and nuisance terms, respectively. *u* stands for the random effect term for phylogenetic relatedness and follows *N*(*0*, *K_pℎylo_σ*^2^). *Z* is the indicator matrix for *u*. *ε* is the residual. Wald test implemented AsREML-R was used to determine the significance of *β*. Because protein abundance can modulate the strength of selection on amino acid composition (i.e., highly abundant proteins make up a larger share of the proteome and are therefore under stronger biosynthetic constraint than less abundant ones^138^), we repeated the association tests on subsets of OGs that are highly abundant in different tissues.

Similarly, for the second hypothesis regarding individual gene activity and environmental transition, we modeled the top three envPCs against five orthogroup-based matrices including presence and absence, premature stop codon occurrence, tip-to-reference dN/dS as well as LLM scores for ESM2^56^ and PlantCAD^55^.

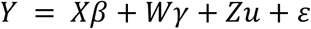

Where *Y* is the envPC, *X* includes all five orthogroup-based gene activity matrices. *W* stands for the nuisance factors including sequencing technology, assembly completeness + duplication level as well as the coverage of the multiple sequence alignment. *β* and *γ* are the fixed effects of predictor terms and nuisance terms, respectively. *u* stands for the random effect term for phylogenetic relatedness and follows 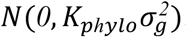 is the indicator matrix for *u*. *ε* is the residual. Where an assembly carried multiple copies of an OG, each copy entered the model as a separate observation; the indicator matrix Z assigns every observation to its assembly so that copies from the same assembly share that assembly’s phylogenetic random effect and are not treated as independent lineages. The per-assembly proportion of multi-copy orthogroups was strongly correlated with assembly duplication (Spearman’s ρ = 0.96) so the duplication covariate captures the genome-wide effect of polyploidy and copy number variation.

Viewing all five OG-based gene activity matrices as non-mutually exclusive mutational consequences on orthologous gene evolution, we tested for their joint effect by contrasting the full model to a reduced model: *Y* = *Wγ* + *Zu* + *ε*. As implemented by R/asremlPlus, the full likelihoods, evaluated using REML estimates, are used in the likelihood ratio tests.

For a subset of high-confidence environmentally associated OGs of particular interest, we tested per-residue association to evaluate the importance of certain residues under the same framework.

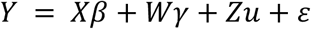

Where *Y* is the envPC, *X* is binarized amino acid allele state (major and others). *W* stands for the nuisance factors including sequencing technology, assembly completeness + duplication level as well as the coverage of the multiple sequence alignment. *β* and *γ* are the fixed effects of predictor term and nuisance terms, respectively. *u* stands for the random effect term for phylogenetic relatedness and follows 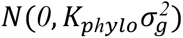 is the indicator matrix for *u*. *ε* is the residual. Wald test implemented AsREML-R was used to determine the significance of *β* in these tests.

We included a phylogenetic permulation process to obtain empirical p-values for all above tests to better control the false discovery rate as described previously ^65^.

#### Molecular evolution

In addition to the phylogenetic mixed model, we also analyzed our data using a state-of-art codon-substitution method, RELAX^76^, to test for shifts in selection intensity across predefined lineages in a given gene tree. Given the computational load (avg. ∼4hr per OG per branch designation on a single thread), we only applied this analysis to the OG showing significant signals in the phylogenetic mixed model and treated the test result as orthogonal evidence to call high-confidence candidates. For each envPC, we labeled the transition branches to both upper and lower 30% of the distribution respectively in a given gene tree and performed two separate tests per gene tree.

#### Curation of abiotic stress responsive genes

Under the assumption that the genes exhibiting expression changes in response to different abiotic stresses are involved in adaptive pathways, we curated such gene lists from previous publications as another orthogonal evidence for high-confidence environmentally associated OG identification. We searched for transcriptomic and proteomic studies associated with keywords: “cold/freezing”, “heat/hot”, “drought/water deficit” and “waterlogging/submergence/flood” in “grasses/Poaceae” species and identified 45 stress-species-study combinations (9-15 species-study combinations per stress category) (Supplementary Table 6)^35,139–166^. For each study, we adopted the list of stress-responsive genes reported by the original authors, applying their published significance criteria without re-analysis. Reported genes were mapped to our OGs via miniProt alignment to the corresponding reference genomes of the original studies. For each stress category, we considered an OG as stress-responsive only if it was reported in at least three species-study combinations. These curated abiotic stress responsive genes were used for enrichment analysis for phylogenetic mixed model candidates as well as filter criteria for high-confidence environmentally associated OGs.

#### Gene Ontology (GO) enrichment analysis

To understand the potential adaptive pathways and biological processes, we performed Gene Ontology enrichment analysis using the “elim” and “classic” algorithm in R/topGO package (v2.50.0)^167^. Maize homologs of the candidate OGs were extracted and maize v5 GO annotation^168^ was used in this analysis. Additionally, we *de novo* annotated GO terms to the reconstructed ancestral OG sequences using deepGO^169^ and performed an enrichment based on this annotation independently.

#### Candidate sequence analysis

For a subset of high-confidence environmentally associated OGs of particular interest, we utilized targetP 2.0^170^, InterProScan^171^ and alphafold3^172^ to further study the mutational effects on signal/transit peptides, protein domains and folding.

### Quantification and Statistical analysis

All details of the statistics applied in this paper are provided in the corresponding method details subsections and figure legends. Statistical tests were performed in R 4.2 environment with designated packages as described.

#### Key resources table

| RESOURCE | SOURCE | IDENTIFIER |
|---|---|---|
| Deposited data |  |  |
| 430 short read Andropogoneae assemblies | This study | https://figshare.com/s/eb35a1d65e4eb6be7c79 |
| 33 Long read Andropogoneae assemblies | Stitzer et al.<sup>53</sup> | PRJEB50280 |
| 207 publicly available genome assemblies | As cited in Hale et al.<sup>49</sup> | As specified in Supplementary Table 4 in this study |
| 57 assemblies generated from publicly available short-read data in the Sequence Read Archive | Hale et al.<sup>49</sup> | https://figshare.com/s/bc19e8f5dae558834cc2 |
| Derived occurrence datasets from BIEN and GBIF and the associated environmental data | This study | https://zenodo.org/records/14968186 |
| Maize proteome dataset | Walley et al.<sup>127</sup> | TableS2 of the original article |
| PlantCAD model weights | Zhai et al.<sup>55</sup> | kuleshov-group/PlantCaduceus_l32 |
| ESM2 model weights | Lin et al. <sup>56</sup> | facebook/esm2_t33_650M_UR50D |
| Arabidopsis curated mutation dataset | Kono et al.<sup>44</sup> | Table S2 of the original article |
| Maize curated mutation dataset | Wang et al.<sup>75</sup> | Table S22 of the original article |
| *a priori* abiotic stress responsive gene list | Ref.<sup>35, 139-166</sup> | As specified in Supplementary Table 6 in this study |
| Software and algorithms |  |  |
| Helixer v0.3.2 | Holst et al.<sup>122</sup> | https://github.com/usadellab/Helixer |
| OrthoFinder v2.5.4 | Emms and Kelly<sup>123</sup> | https://github.com/davidemms/orthofinder |
| R/phangorn package v2.12.1 | Schliep<sup>124</sup> | https://cran.r-project.org/web/packages/phangorn/index.html |
| miniProt v0.13.0 | Li<sup>125</sup> | https://github.com/lh3/miniprot |
| MAFFT v7.520 | Katoh and Standley<sup>126</sup> | https://mafft.cbrc.jp/alignment/server/index.html |
| RAxML v8.2.12 | Stamatakis<sup>129</sup> | https://github.com/stamatak/standard-RaxML |
| ASTRAL-Pro2 | Zhang and Mirarab<sup>130</sup> | https://github.com/chaoszhang/A-pro |
| R/ape v5.8-1 | Paradis et al.<sup>173</sup> | https://cran.r-project.org/web/packages/ape/index.html |
| R/canprot v2.0.0 | Dick<sup>174</sup> | https://github.com/jedick/canprot |
| SeqKit v0.15.0 | Shen et al.<sup>133</sup> | https://github.com/shenwei356/seqkit |
| R/MSA2dist v1.2.0 | Ullrich<sup>175</sup> | https://github.com/kullrich/MSA2dist |
| ASREML-R v4.2.0 | Butler et al.<sup>135</sup> | https://asreml.kb.vsni.co.uk/knowledge-base/release-notes-asreml-r-4-2/ |
| R/phylolm v2.6.5 | Ho and Ané<sup>137</sup> | https://cran.r-project.org/web/packages/phylolm/index.html |
| RELAX (as implemented in HyPhy 2.5.49) | Wertheim et al.<sup>76</sup> | https://github.com/veg/hyphy-analyses |
| R/topGO v2.50.0 | Alexa et al.<sup>167</sup> | https://bioconductor.org/packages/release/bioc/html/topGO.html |
| deepGO2 | Tawfiq et al.<sup>169</sup> | https://github.com/bio-ontology-research-group/deepgo2 |
| targetP 2.0 | Almargo Armenteros et al.<sup>170</sup> | https://github.com/JJAlmagro/TargetP-2.0 |
| InterProScan5 | Jones et al. 2014<sup>171</sup> | https://github.com/ebi-pf-team/interproscan |
| alphafold3 | Abramson et al.<sup>172</sup> | https://github.com/google-deepmind/alphafold3 |

## Supplemental Information

Document S1. Supplementary Figures S1-S10

Supplementary Table 1. Power estimates based on the simulation results

Supplementary Table 2. Phylogenetic mixed model results

Supplementary Table 3. Gene ontology enrichment results

Supplementary Table 4. Metadata of genome assemblies analyzed in this study

Supplementary Table 5. The reconstructed ancestral amino acid sequences of the 22,363 OGs

Supplementary Table 6. The list of studies on grass abiotic stress expression profiles and curated convergent responsive genes

