## Supplementary Figure for "Convergent genome- and gene-level constraints shape repeated environmental adaptation in grasses"

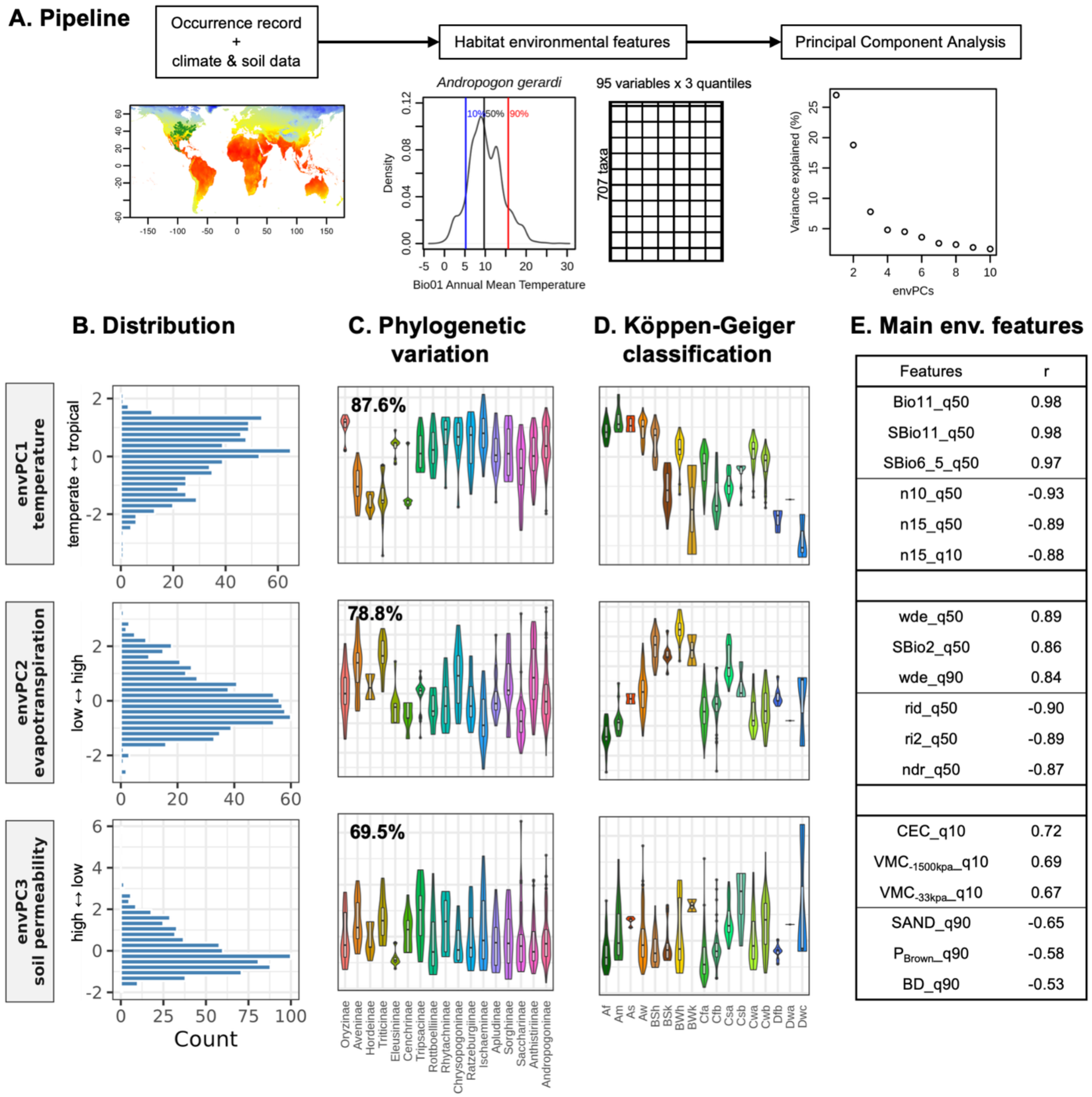


**Supplementary Figure S1. Species habitat environmental characterization.**

A. Flowchart of the pipeline. Occurrence records of any given species is retrieved from both GBIF and BIEN databases. The environmental variables associated with the coordinates were retrieved. For each variable, 10, 50, 90 percentiles of the distribution were calculated and used to derive the environmental principal components (envPCs) that represent the environmental niches for a given species. B. The distributions of each envPC. C. The distributions of each envPC, grouped by major subtribes. D. The distributions of each envPC, grouped by the KG3 classification. E. The top environmental variables with the largest loading on each of the envPC.

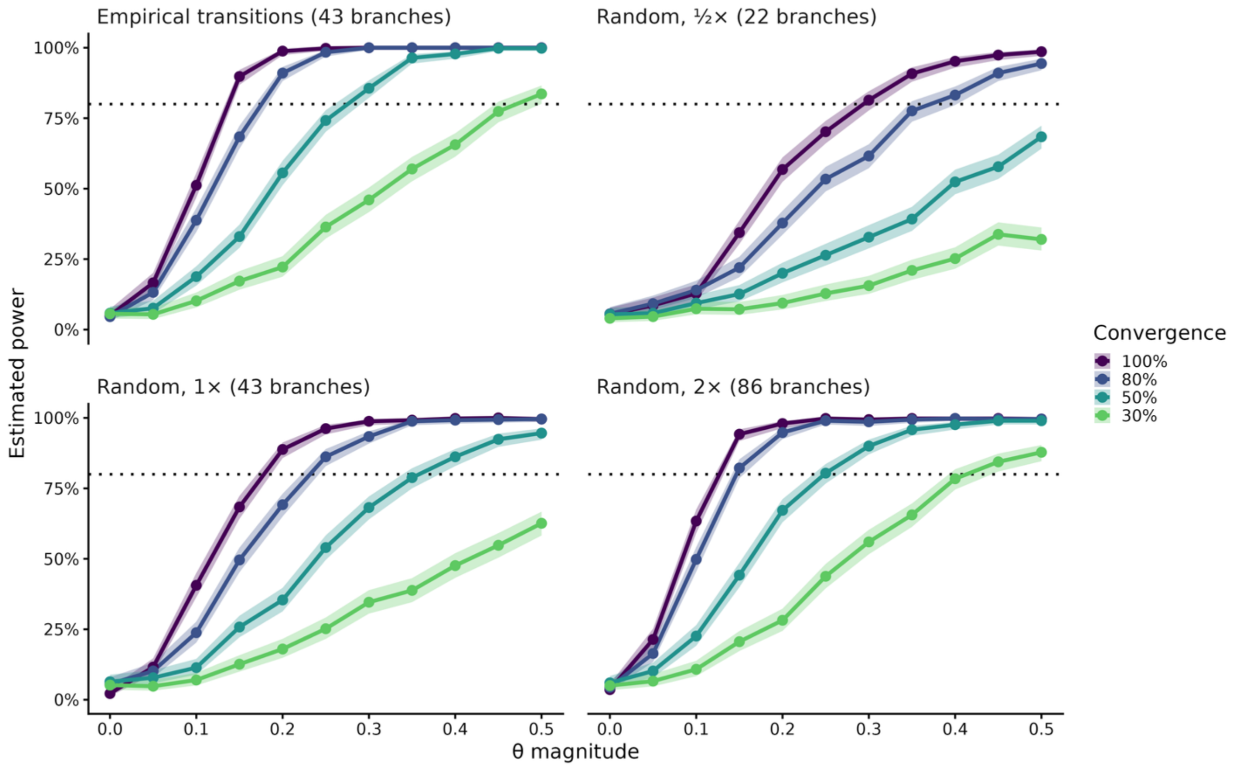


**Supplementary Figure S2. Power curves of phylogenetic mixed model approach under different simulated scenarios.**

We simulated phenotypic (Y) and genotypic variables (X) that follow independent Brownian motion (BM) but share mean shifts at the transition branches in the phylogeny and evaluated the power of the phylogenetic mixed model in detecting such co-shift between the variables. The power curves under different simulated parameter configurations are shown. We tested four transition scenarios: (1) empirical transition for envPC1 as inferred in this study (empirical transitions), random transition of (2) the same (random, 1X), (3) twice (random, 2X) or (4) half (random, 1/2X) the empirical ones. Furthermore, we simulated various co-varying effect sizes (θ) from 0 to 0.5 with a step of 0.05 and different levels of convergent evolution by subsampling the transition branches to 100%, 80%, 50% or 30% of all transitions. The solid dots indicate average power across 500 simulations and shaded area denotes 95% confidence interval. The power of the phylogenetic mixed model increases as the convergence level and relative co-shift size increase.

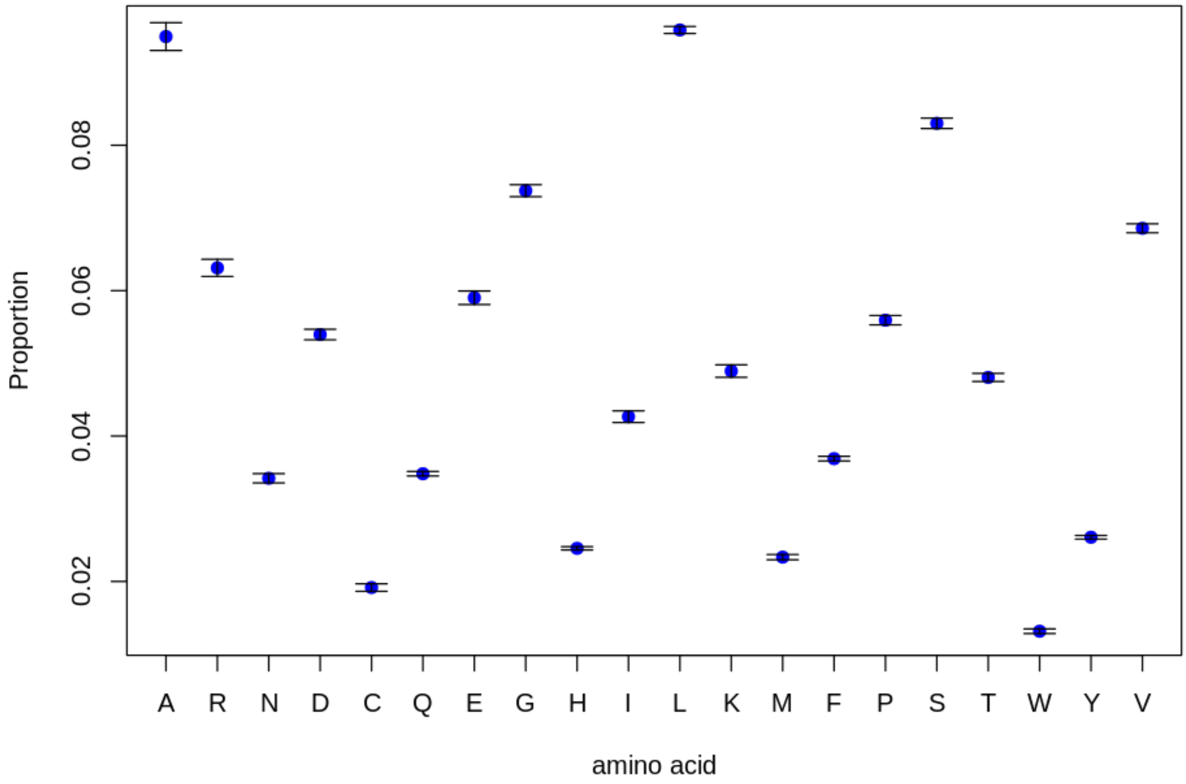


**Supplementary Figure S3. Variation in the amino acid composition among diverse grass species.** The distributions of the proportion of each amino acid in the proteogenic amino acid sequence among the 707 taxa. Mean and one standard deviation is shown.

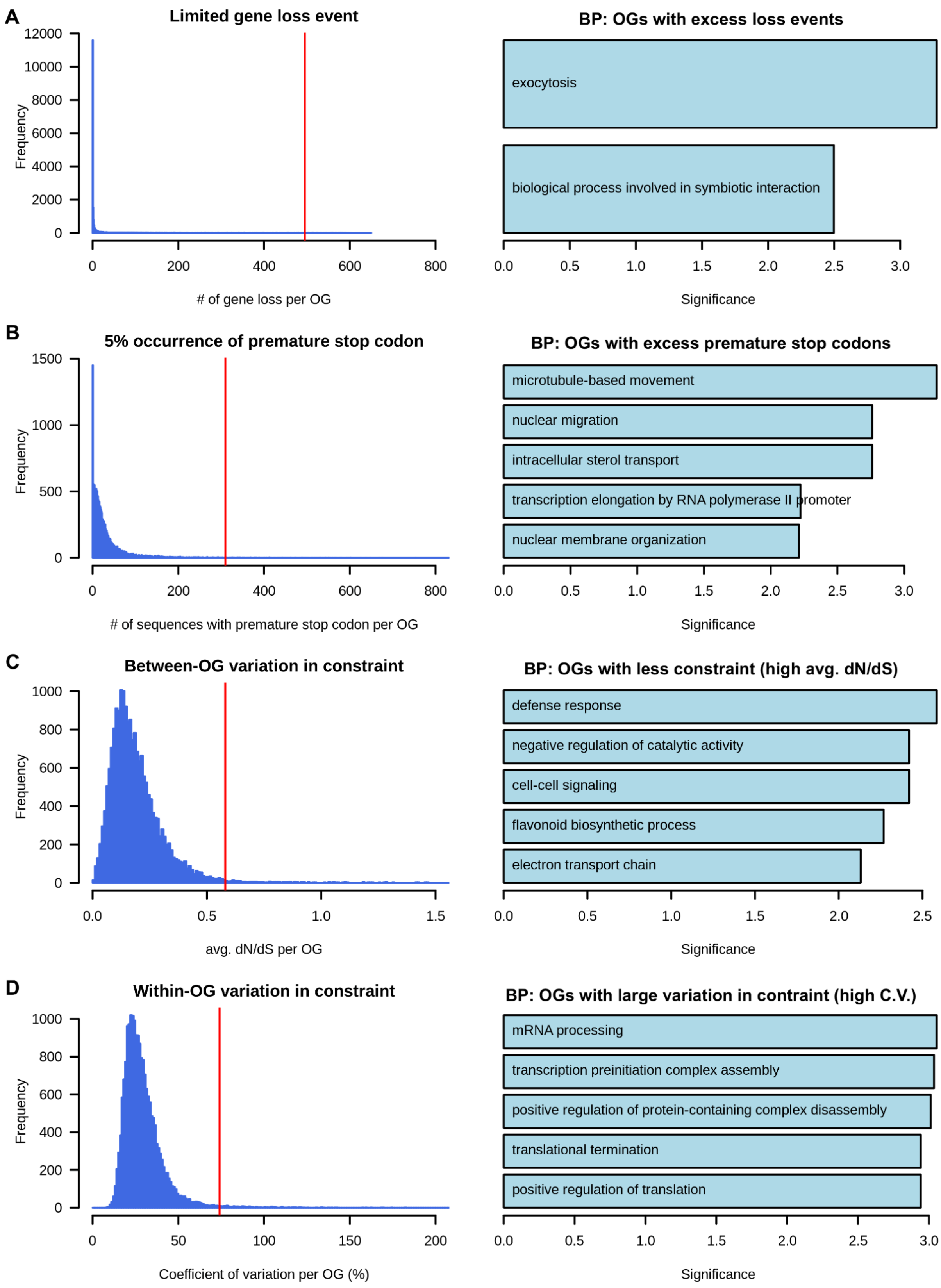

A
Limited gene loss event
Frequency
### of gene loss per OG
BP: OGs with excess loss events
exocytosis
biological process involved in symbiotic interaction
Significance
B
5% occurrence of premature stop codon
### of sequences with premature stop codon per OG
BP: OGs with excess premature stop codons
microtubule-based movement
nuclear migration
intracellular sterol transport
transcription elongation by RNA polymerase II promoter
nuclear membrane organization
C
Between-OG variation in constraint
avg. dN/dS per OG
BP: OGs with less constraint (high avg. dN/dS)
defense response
negative regulation of catalytic activity
cell-cell signaling
flavonoid biosynthetic process
electron transport chain
D
Within-OG variation in constraint
Coefficient of variation per OG (%)
BP: OGs with large variation in contraint (high C.V.)
mRNA processing
transcription preinitiation complex assembly
positive regulation of protein-containing complex disassembly
translational termination
positive regulation of translation

**Supplementary Figure S4. Multiple sequence alignment (MSA) based gene activity inference**
Histograms of the numbers of sequence absence (A), premature stop codon occurrence (B), average dN/dS per OGs (Between-OG variation; C) and coefficient of variation of dN/dS per OG (Within-OG variation; D) are shown. 99 percentiles of the distributions are indicated by the vertical line in red. The right panels show top gene ontology (GO) terms (BP: biological process) enriched by the OGs above 99 percentile of the distributions (i.e., OG with elevated evolutionary rates or divergent selection).

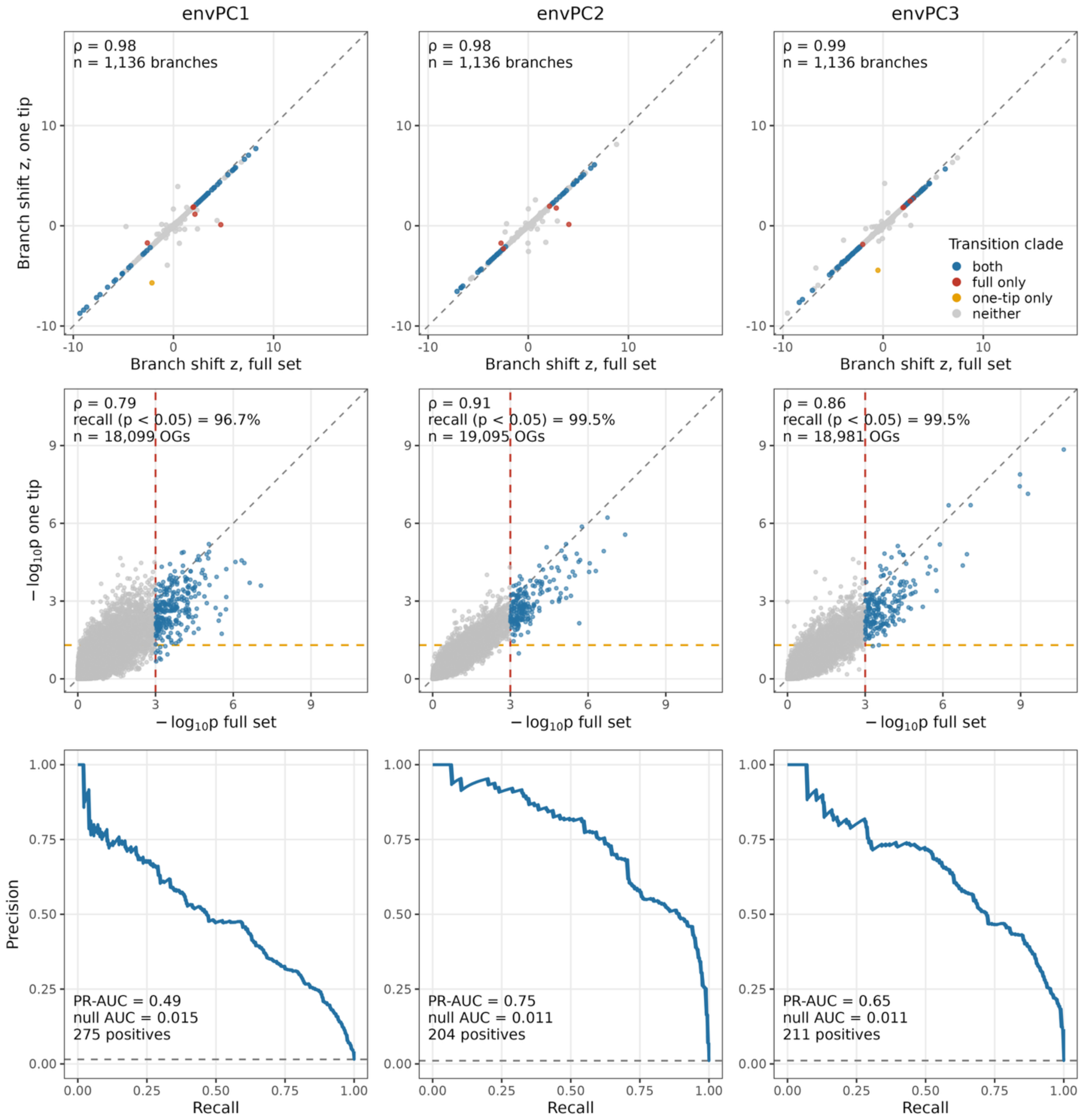


**Supplementary Figure S5. Transition calling and phylogenetic Association results are robust to sampling a single genome per species.**

Both transition calling and phylogenetic association analyses were repeated after subsampling to one representative genome per species ("one tip"). These results are compared with the full 687-genome dataset ("full set") for each of the first three environmental principal components (envPC1-3; columns). ***Top row*** Branch-level environmental shift z-scores from the full set (x-axis) and the one-tip set (y-axis) for the 1,136 branches shared by both trees. Points are colored by whether the branch was called a transition clade in both analyses (blue), in the full set only (red), in the one-tip set only (orange), or in neither (grey). The dashed line marks y = x. ρ, Spearman's rank correlation. ***Middle row*** OG-level association significance (-$\log_{10}$p) from the full set versus the one-tip set (n = 18,099, 19,095 and 18,981 OGs for envPC1-3). The dashed line marks y = x. ρ, Spearman's rank correlation. The vertical red dashed line marks the full-set significance threshold at p of 0.001. The horizontal dashed line marks p = 0.05 for one-tip set results. Orthogroups passing the full-set threshold are shown in blue. Recall is the proportion of these that also reach $p < 0.05$ in the one-tip analysis. ***Bottom row*** Precision–recall curves showing how well one-tip p-values recover the OGs that are significant in the full set (275, 204 and 211 positives for envPC1-3). The horizontal dashed line shows the null expectation, which equals the proportion of positives among all OGs (null AUC = 0.015, 0.011 and 0.011). The PR-AUCs (0.49, 0.75 and 0.65) are 45–68-fold above this null.

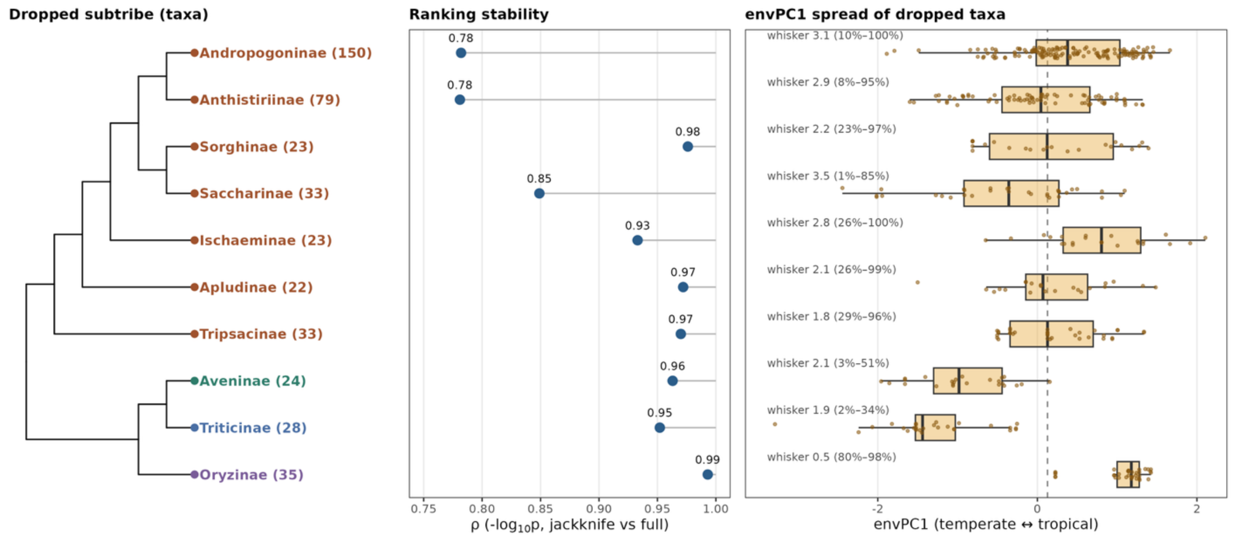


**Supplementary Figure S6. Leave-one-subtribe-out jackknife shows that envPC1 associations are robust to removal of any single lineage.**

Each of ten subtribes with more than 20 taxa was removed in turn and the OG-envPC1 association analysis was re-run. Rows correspond to the subtribe dropped in each replicate. ***Left*** Cladogram of the dropped subtribes, with the number of taxa removed in parentheses. Colors mark tribe: Andropogoneae (brown), Poeae (green), Triticeae (blue) and Oryzeae (purple). ***Middle*** Ranking stability, measured as Spearman's ρ between OG $-\log_{10}$ p from each jackknife replicate and from the full dataset. ρ ranged from 0.78 to 0.99 (median 0.955). The lowest values came from removing the two largest subtribes, Andropogoninae (150 taxa; ρ = 0.78) and Anthistiriinae (79 taxa; ρ = 0.78). ***Right*** envPC1 values of the taxa removed in each replicate. Boxes show the median and interquartile range, whiskers extend to 1.5 × IQR, and points show individual taxa. Each label gives the envPC1 whisker length of the dropped taxa, followed by the percentile interval of the full-dataset envPC1 distribution that they span. The dashed line marks the full-dataset median.

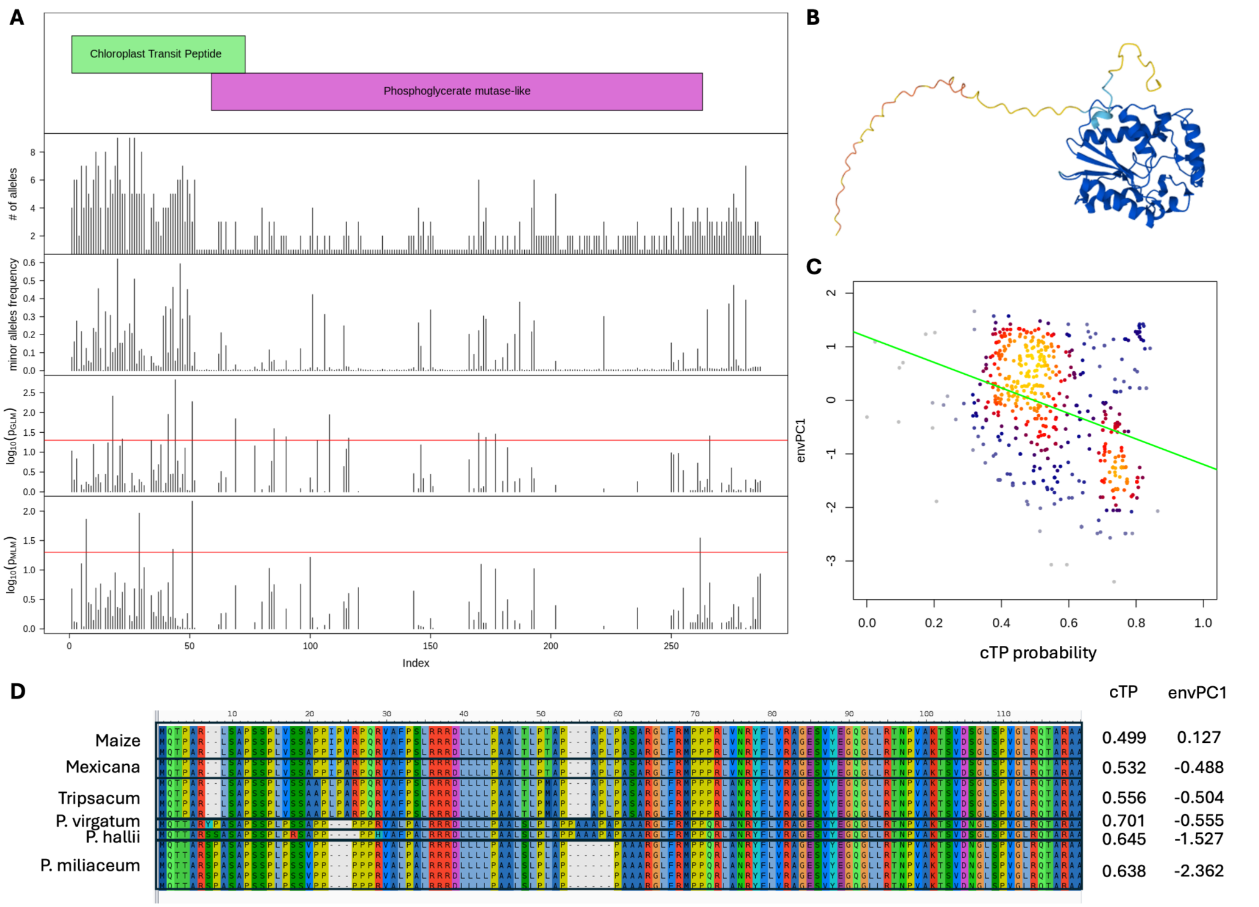


**Supplementary Figure S7. Sequence analysis of OG0018915.**

A. Protein domains as predicted by InterProScan and targetP 2.0. The numbers of unique amino acids, the frequency of non-consensus alleles and the residue-based association of envPC1 with and without accounting for phylogenetic relatedness are shown. Diversity and significant environmental association are concentrated in the predicted chloroplast transit peptide (cTP) region. B. Predicted protein structure of the protein. C. The predicted probability of cTP presence is significantly associated with envPC1 (negatively correlated). D. Multiple sequence alignment of the predicted cTP region among selected Panicoideae sequences.

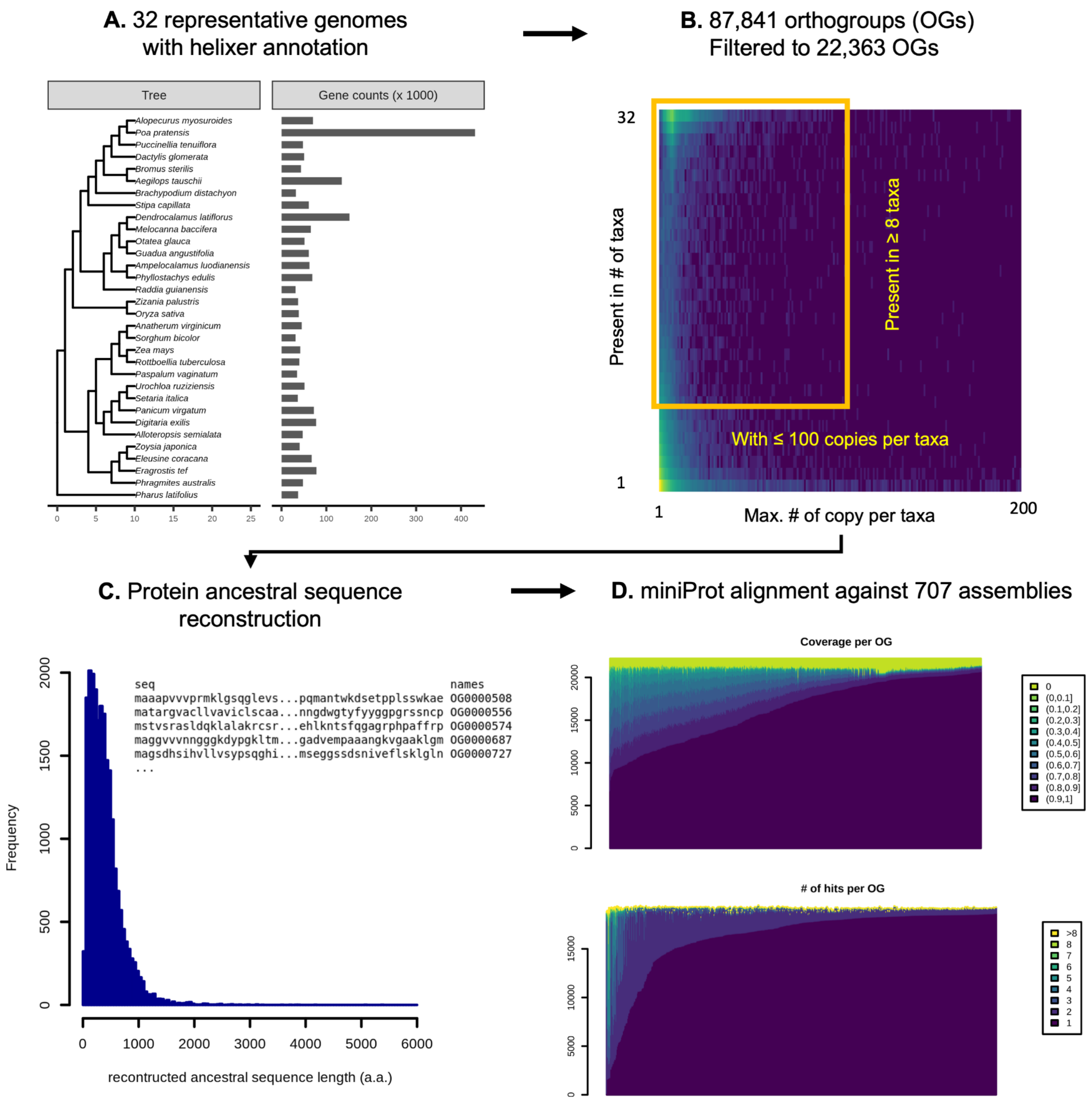


**Supplementary Figure S8. Orthogroup construction and genomic alignment.**
**A**. The phylogeny and the helixer-annotated gene counts of the 32 representative Poaceae genomes. The gene counts match the ploidy expectation of the taxa. **B**. Using orthofinder, we constructed 87,841 orthogroups in total. Requiring the orthogroup to be present in at least 8 of the 32 representative genomes and with no more than 100 copies per taxon filtered to 22,363 orthogroups. **C**. we reconstructed the ancestral amino acid sequences for each orthogroup using R/phangorn and the length distribution of the reconstructed proteins is shown. D. The reconstructed protein sequences were queried against all 707 assemblies in this study using miniProt. In most assemblies, we got at least one hit with >50% alignment coverage per orthogroup. The variation in alignment coverage was accounted for in the phylogenetic mixed linear model.

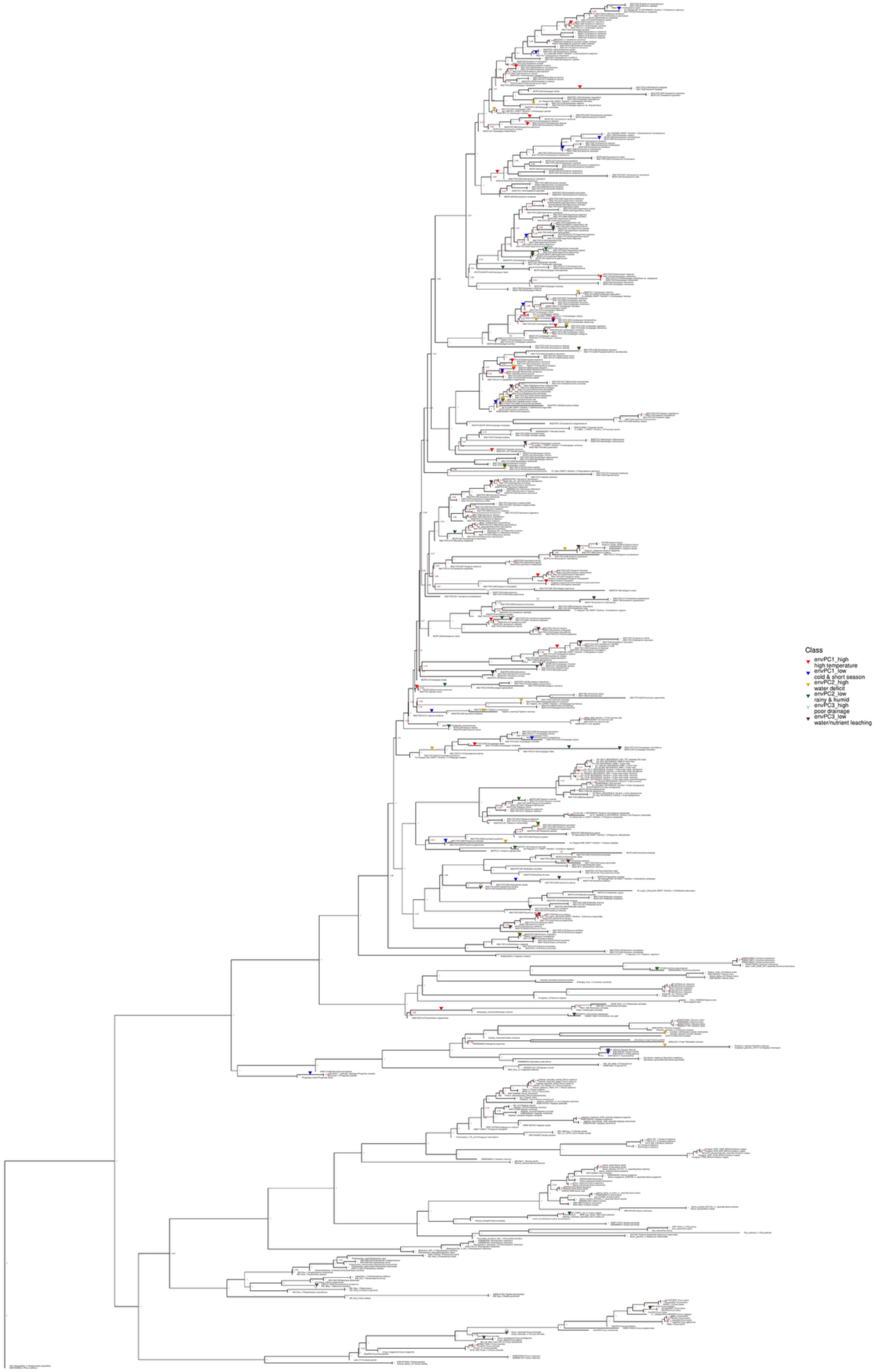
Class
envPC1_high
high temperature
envPC1_low
cold & short season
envPC2_high
water deficit
envPC2_low
rainy & humid
envPC3_high
poor drainage
envPC3_low
water/nutrient leaching

**Supplementary Figure S9. The phylogeny of the 687 taxa analyzed in this study.**

We generated the phylogenetic tree based on the gene trees of the orthogroups containing the angiosperm353 loci using ASTRAL-Pro2. The tip labels are in the format of assemblyID:species_name. The red numbers at nodes indicate the branch supports measured as local posterior probabilities. We indicated the branches where transitions to various environments occurred.

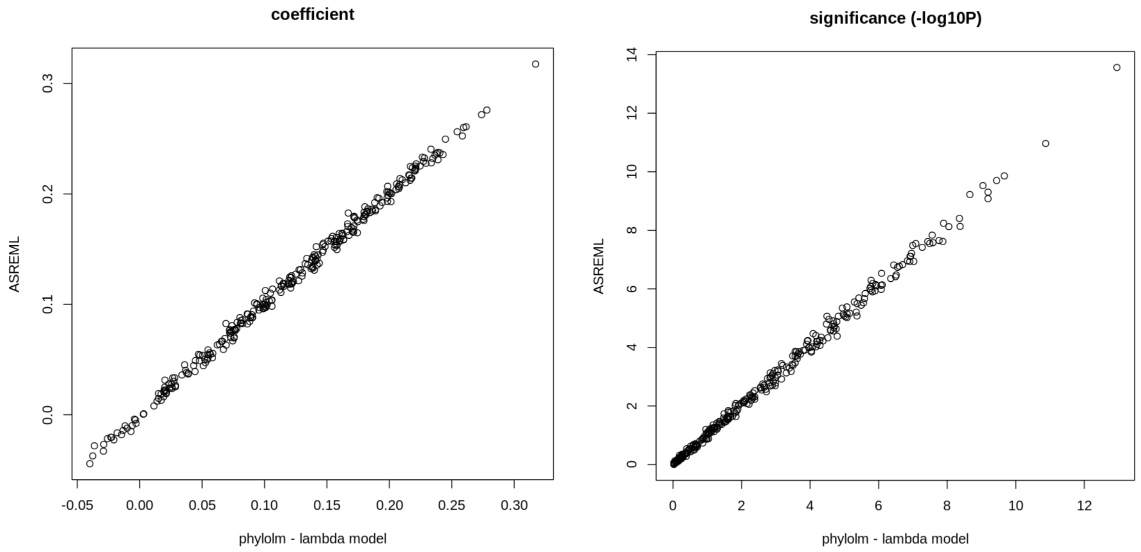


**Supplementary Figure S10. Comparison between ASREML-R and phylolm**

Based on the empirical phylogeny in this study, 300 simulated trait-pairs of different association levels were used to compare the performance of ASREML-R and phylolm in detecting phylogenetic co-variation. There is high correlation between the two methods for both coefficient estimates and significance (Spearman's rho > 0.998).
